# Substantial cold tolerance in all life stages of the biting midge, *Culicoides nubeculosus* (Diptera: Ceratopogonidae)

**DOI:** 10.64898/2025.12.04.692099

**Authors:** Lucy M. Devlin, Ross N. Cuthbert, Melanie Nicholls, Archie K. Murchie, Connor G. G. Bamford, Jaimie T. A. Dick, Eric R. Morgan, Son T. Mai, Marion England

## Abstract

In temperate regions, vector-borne disease risk is mediated by cold winter conditions, however, the cold tolerance of key vector taxa remains poorly understood. *Culicoides* biting midges are the primary vectors of several pathogens of medical and veterinary importance including bluetongue virus, where seasonal cold weather in temperate regions limits midge activity and pathogen transmission. Here, we provide the first comprehensive assessment of cold tolerance across all developmental stages of *Culicoides nubeculosus*, a widely used laboratory species that is endemic to northern Europe. Eggs, first-instar larvae, fourth-instar larvae, pupae, and adults were exposed to acute (1 h) and extended (6 and 24 h) cold treatments spanning −1 to −18 °C, with survival, development, emergence, and adult wing size quantified*. Culicoides nubeculosus* showed substantial but stage-specific cold tolerance, with survival limits of ≤ −18 °C for eggs, −14 °C for pupae, −10 °C for L1 larvae and adults, and −7 °C for L4 larvae. While the effect of cold exposure duration varied across temperatures and life stages, extended exposure generally reduced survival at lower temperatures. Cold stress caused sublethal effects, including reduced adult emergence when eggs or larvae were exposed and reductions in adult wing size of up to ∼10%, depending on the life stage. These results reveal substantial cold tolerance across the full life history of *C. nubeculosus*, suggesting that factors beyond temperature influence population phenology. Our findings provide new insights into *Culicoides* ecology, with implications for seasonal vector population dynamics and arbovirus transmission risk in temperate regions.

**Graphical Abstract:** 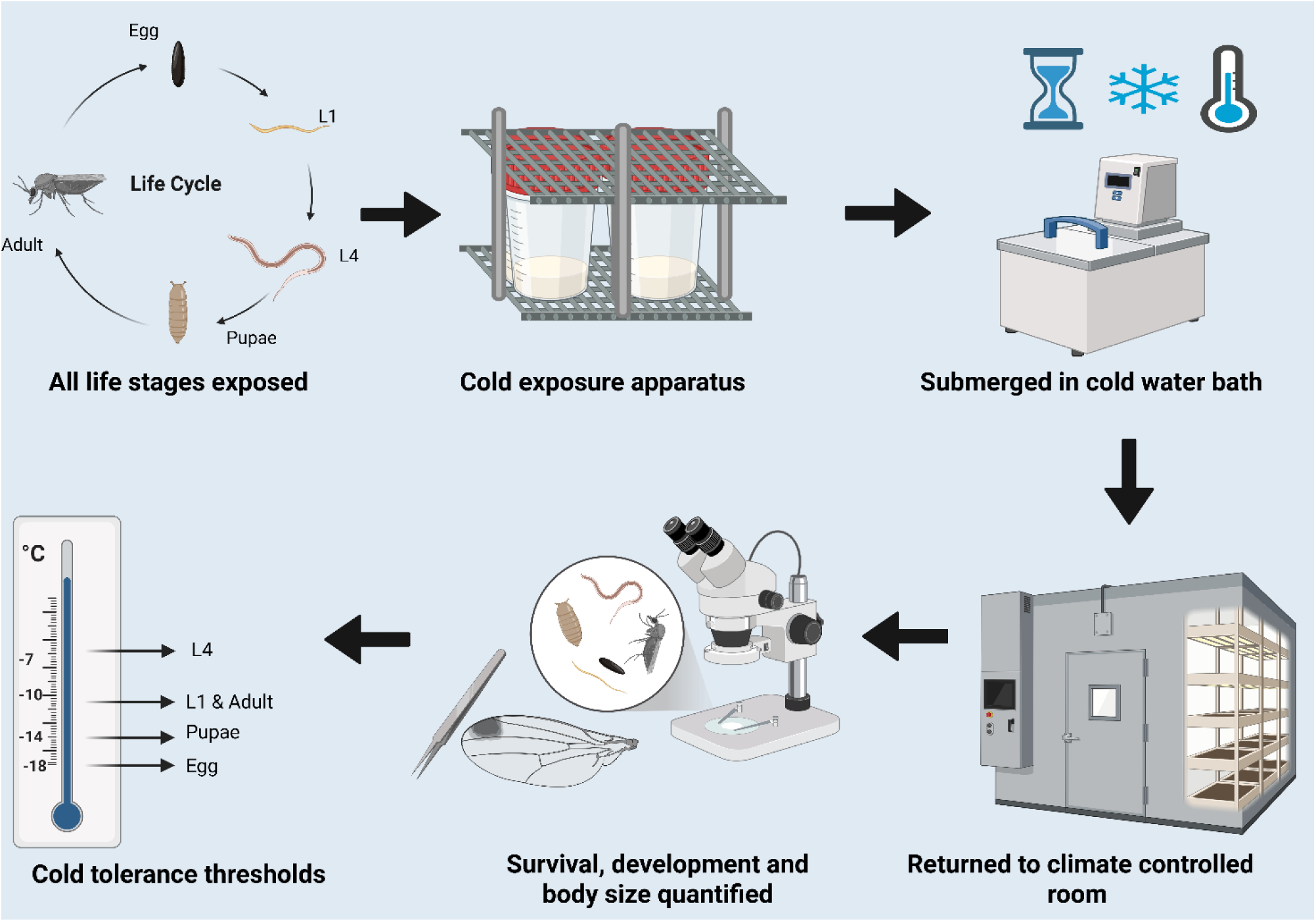

## 1. Introduction

Vector-borne diseases (VBDs) represent a substantial threat to global health, agriculture, and economic stability, with environmental changes aggravating disease risks and promoting their emergence and re-emergence in various regions (Gubler 2010, Chala and Hamde 2021). Key arthropod vectors include mosquitoes, ticks, and biting midges (*Culicoides* spp.), with certain species capable of transmitting a wide range of pathogens and parasites to humans, livestock, and wildlife (Collins et al. 2018, Socha et al. 2022). Climate change is a key driver of shifting VBD dynamics, influencing both the geographic spread and emergence of vectors and associated disease. Global temperatures have risen markedly in recent decades, and climate models project continued warming throughout the 21st century, together with an intensification of climatic extremes such as extended warm spells and sudden cold snaps (Caminade et al. 2019, Ryan et al. 2019, Chala and Hamde 2021, Wu et al. 2025). Such climatic shifts are important because arthropod vectors exhibit strong temperature dependence in traits linked to their survival, reproduction, distribution, and ultimately the duration of transmission seasons. While warming trends often extend the active transmission season, cold winter conditions and sudden cold events can still constrain vector persistence, particularly in temperate regions (Carpenter et al. 2013, Caminade et al. 2019, Sanders et al. 2019, Chapman et al. 2020, Chala and Hamde 2021).

*Culicoides* biting midges (Diptera: Ceratopogonidae) are small, hematophagous insects with complex life cycles and diverse ecological requirements (Carpenter et al. 2013, Purse et al. 2015). They occupy a wide range of habitats, and several species are strongly associated with anthropogenic substrates, such as livestock dung, slurry, and manure-enriched soils, which can alter local microclimates and support high midge populations that increase the likelihood of disease transmission among host animals (Uslu and Dik 2010, Thompson et al. 2013, Purse et al. 2015, Werner et al. 2020). The life cycle of *Culicoides* includes an egg stage, four larval instars (L1–L4), a pupal stage, and the adult (Carpenter et al. 2013, Purse et al. 2015). Each developmental stage has specific ecological requirements and is strongly influenced by environmental factors such as temperature and moisture (Purse et al. 2015, Sanders et al. 2019). Eggs are laid in moist substrates, and larvae develop in semi-aquatic habitats, typically making up the longest phase of the life cycle. Pupation occurs near the substrate surface, followed by the emergence of adults. Adults are short-lived, generally surviving a few days to several weeks depending on temperature and humidity, and are mainly crepuscular (Purse et al. 2015, Steyn et al. 2016). These midges serve as primary vectors of several economically important arboviruses, including bluetongue virus (BTV), African horse sickness virus (AHSV) and Schmallenberg virus (SBV) (Carpenter et al. 2013, Purse et al. 2015, Collins et al. 2018, England et al. 2020). While *Culicoides* are best known as vectors of arboviruses, emerging evidence indicates they may also transmit non-viral pathogens such as *Leishmania* (Modrý et al. 2025). Of the approximately 1,400 described species globally, a small proportion of *Culicoides* are confirmed vectors, due to both variation in vector competence and limited experimental testing (Carpenter et al. 2013, Purse et al. 2015).

Higher temperatures can accelerate *Culicoides* larval development and virus replication but may also reduce adult lifespan and disrupt transmission dynamics once optimal thermal limits are exceeded (Rozo-Lopez et al. 2022, Banerjee et al. 2024). In contrast, exposure to low temperatures poses considerable challenges to survival and slows development across life stages, particularly in temperate regions where seasonal cold can limit persistence (Van Den Eynde et al. 2021). Overwintering of *Culicoides* and their arboviruses remains poorly understood, particularly in temperate regions, where the mechanisms underlying cold survival are not well characterised (Carpenter et al. 2013, White et al. 2017, Sanders et al. 2019, Sick et al. 2019). This has been most apparent with the recent outbreak of bluetongue virus serotype 3 (BTV-3) in northern Europe, which has successfully overwintered, after initially emerging in 2023 (Dijkstra et al. 2025, King et al. 2025). However, several mechanisms have been proposed to explain persistence of both *Culicoides* and the viruses they transmit, involving adult survival during mild periods and cold-tolerant immature stages such as eggs and late-instar larvae (White et al. 2005, Lysyk and Danyk 2007, Steyn et al. 2016, McDermott et al. 2017). Eggs in particular appear to have the highest cold tolerance, possibly due to their desiccation resistance and small size, while larvae and pupae tend to be more susceptible to cold stress (McDermott et al. 2017). Despite this, evidence suggests that third and fourth instar larvae can persist through winter under certain conditions, even in partially frozen habitats (Lysyk and Danyk 2007). However, little is known about the relative cold tolerance of different life stages in *Culicoides*, including variation among larval instars, adult survival capacity, and tolerance across different durations of cold exposure (McDermott et al. 2017). In particular, there is an absence of these studies across northern European *Culicoides* vectors. Addressing these knowledge gaps is critical, as differences in cold tolerance across life stages may determine whether infected *Culicoides* populations can persist through winter, thereby sustaining arbovirus transmission across seasons (Steyn et al. 2016, Sanders et al. 2019).

This study thus investigates the cold tolerance of *Culicoides nubeculosus* (Meigen), a temperate species native to northern Europe that has been maintained in colony and, being refractory to major arboviruses, provides a safe and widely used laboratory model for behavioural studies (Boorman 1974, Carpenter et al. 2013). Based on previous findings in *Culicoides sonorensis* Wirth and Jones (McDermott et al. 2017), we hypothesised that (i) eggs would exhibit the greatest cold tolerance, (ii) larvae would be the least tolerant, (iii) longer durations of cold exposure would result in progressively higher mortality across all stages, and (iv) sublethal cold stress would reduce subsequent development rates and adult body size. We thus examined the responses of eggs, early (L1) and late (L4) instar larvae, pupae, and adults of *C. nubeculosus* to cooling regimes and cold exposure durations. For each stage, we quantified immediate survival and subsequent development, as well as trait-mediated effects of cold exposure through morphometric analysis of midge wing sizes. By considering all stages of the life cycle, this study provides the first comprehensive assessment of cold tolerance in a northern European species of *Culicoides*, offering insights into their overwintering ecology.

## 2. Methods

### 2.1 Study organisms

All life stages of *C. nubeculosus* were obtained from a long-established colony maintained at The Pirbright Institute, Woking, United Kingdom. The colony was originally established in 1969 from field-collected adults in Hertfordshire, United Kingdom (Bellekom et al. 2023), and has been maintained as a closed colony derived from those founding individuals since then

(Boorman 1974). The colony is maintained under controlled environmental conditions: 27°C, 55±10% relative humidity, and a 12:12 hour light: dark photoperiod. Females are blood-fed three times per week on donor defibrinated horse blood (TCS Biosciences Ltd.) using a Hemotek membrane feeding system, and egg papers are collected on each feeding day, representing oviposition from the previous blood meal. For the purposes of our study, eggs were collected two days after blood feeding, and first-instar (L1) larvae were collected one day after hatching. Fourth instar (L4) larvae and pupae were collected from the oldest rearing trays and L4 were identified based on larval size. Pupae were approximately two days old at the time of collection. Adults were collected one day after emergence.

### 2.2 Experimental setup

All temperature exposure experiments were conducted using a Julabo refrigerated water bath

(Julabo MA F12, UniGreenScheme, UK) filled with Thermal G bath fluid (GPE Scientific Ltd, UK), allowing for precise control of low temperatures down to −18°C. A custom fabricated stainless-steel metal cage was used to hold four upright 125ml specimen pots (polypropylene container, Scientific Laboratory Supplies, UK), ensuring they were fully submerged without direct contact with the bath walls or heating/cooling elements (Figure 1). This cage design minimised unintended temperature gradients and allowed uniform water circulation around all samples. Internal temperatures were verified using a calibrated precision thermometer (RS 1720 Wired Digital Thermometer, RS Components, UK) placed inside a representative pot to confirm alignment with the bath’s digital settings. Control treatments were conducted using the same pot and cage setup in the water bath, but with the bath switched off, exposing samples to ambient laboratory temperature (approximately 20 ± 2°C), thereby ensuring all conditions aside from thermal exposure were identical.

**Figure 1.**
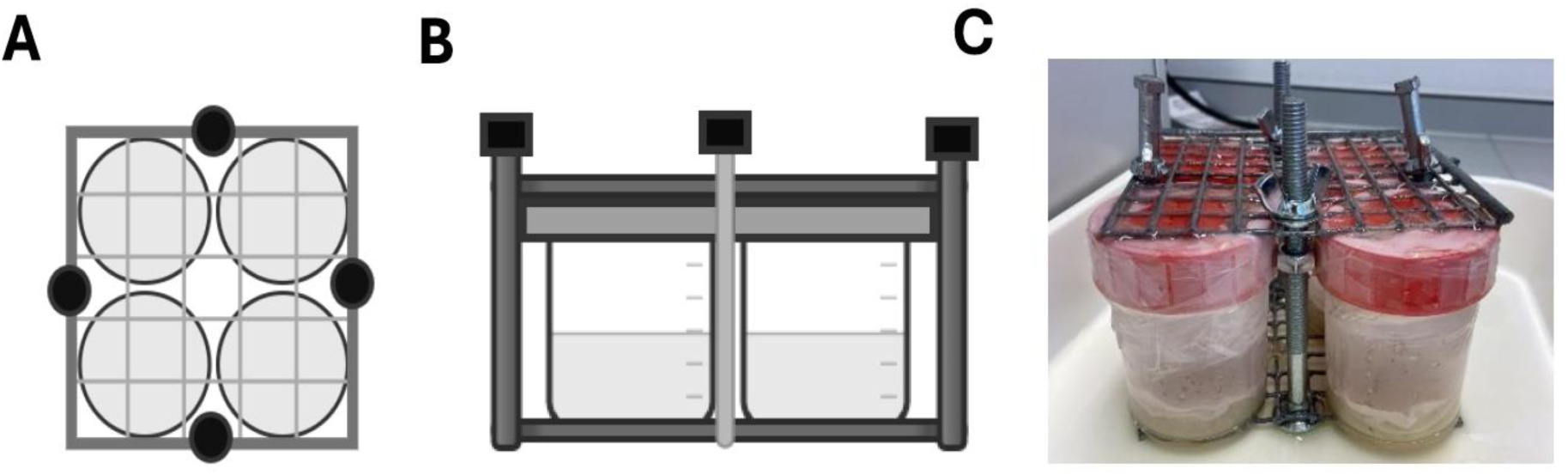
Cold exposure apparatus. Diagram showing the metal cage from above (A) and from the side (B) holding four 125 ml pots, used to ensure uniform temperature exposure across all replicates. The photograph (C) illustrates the actual setup used during experiments.

**Alt text:** Graphs showing the percentage of *Culicoides nubeculosus* L4 larvae surviving at 1 hour and 10 days post cold exposure to temperatures of −1 °C, −6 °C, −7 °C, and control for durations of 1, 6 and 24 hours.

### 2.3 Container setup and substrates

The container setup and substrate composition (summarised in Supplementary Table S1), defined here as the physical materials and media used to support each life stage (e.g., polyester wadding, rearing fluid), are described below (Section 2.4). Each life stage required slightly different rearing setups to support survival, which were established based on prior experience maintaining the colony. The rearing fluid, consisting of dechlorinated tap water mixed with Nutrient Broth No. 2 (ThermoFisher Scientific), dried grass meal (Emerald Green Feeds, Poucher & Sons Ltd, UK), and wheatgerm (The Easy Health Store Limited), was collected from the main colony trays and carefully sieved through fine mesh cloth to remove any unintended life stages or debris before use. All setups were prepared on the day of each trial, with submersion into the water bath occurring approximately one hour after preparation.

### 2.4 Experimental protocols

#### 2.4.1 Life stage specific protocols

For eggs, hatching was monitored across several time points (days 4, 8, 12, and 14 postexposure) to capture temporal variation in emergence, but overall survival was determined at the final time point (day 28; Table 1). Early-instar larvae (L1) were assessed only at day 28 due to their small size and fragility. Fourth-instar larvae (L4) and adults were assessed one hour post-exposure and again following a recovery period under colony conditions (day 10 for L4; day 3 for adults). Pupae were assessed only at day 6, when adult emergence could be reliably determined. Each treatment condition was replicated four times. For each treatment condition, four replicate pots containing 25 individuals were prepared (N = 100 per life stage per treatment).

**Table 1.** Observation timepoints and measured outcomes for each life stage of *Culicoides nubeculosus*. Individuals were monitored at fixed timepoints to assess hatching success, survival, and developmental stage. At the final observation, individuals were categorised as subadults or adults, depending on their life stage.

| Life stage | Observation timepoints | Measured outcomes |
| --- | --- | --- |
| Eggs | Day 4, day 8, day 12, day 14, day 28 | Hatching success (days 4–14); subadult and adult counts and survival (day 28) |
| Early instar larvae (L1) | Day 28 | Survival (day 28); subadult and adult counts (day 28) |
| Late instar larvae (L4) | Hour 1; day 10 | Survival (hour 1 and day 10); subadult and adult counts (day 10) |
| Pupae | Day 6 | Adult emergence (day 6) |
| Adults | Hour 1; day 3 | Survival (hour 1 and day 3) |

#### 2.4.2 Eggs

Groups of 100 eggs (approximately one day old) were collected using a fine paintbrush and divided evenly among four replicate Petri dishes (25 eggs per dish). Each dish contained filter paper placed over a circle of polyester wadding dampened with 3 ml of rearing fluid in a 35 mm Petri dish. The dishes were sealed with Parafilm, placed into 125 ml plastic pots (as described above), secured within the metal cage, and submerged in the water bath. After cold exposure, the egg papers from each Petri dish were transferred to fresh 30 ml plastic pots (4.5 × 4.5 × 3 cm; Ambican UK Ltd) containing 2 ml of sieved colony rearing fluid to promote hatching. After 14 days, the entire contents of the 30 ml pots were transferred to 125 ml pots covered with mesh lids to allow for further development of the larvae that had hatched from the eggs. Hatch rates were recorded using a microscope at days 4, 8, 12, and 14 postexposure. To support larval development, 0.2 g of dried grass meal was added on day 2, and 1 ml of sieved colony rearing fluid was added every two days. Final counts at day 28 recorded numbers progressing to subadult and adult stages (Table 1).

#### 2.4.3 Early instar larvae (L1)

Instar 1 larvae (L1) (∼1 days post-hatching) were collected using a micropipette under a microscope and transferred into 125 ml pots containing polyester wadding and 15 ml sieved colony water. After cold exposure, pots were returned to colony conditions. Survival was assessed at day 28 post-exposure by counting individuals that developed into subadult or adult stages (Table 1). Grass meal (0.2 g) was provided on the second day, and 2 ml of colony water was provided every two days.

#### 2.4.4 Late instar larvae (L4)

Instar 4 larvae (L4) were identified visually from colony trays and transferred directly into 125 ml pots containing polyester wadding and 15 ml of sieved colony water. After cold exposure, initial survival was assessed 1 hour post-exposure by observing movement or response to gentle probing with a pipette (McDermott et al. 2017). Surviving individuals were then maintained in colony conditions, and survival and developmental outcomes were recorded again on day 10 (Table 1).

#### 2.4.5 Pupae

Pupae were transferred using a fine paintbrush into prepared pots containing polyester wadding, 12 ml of sieved colony rearing fluid, and a filter paper layer prior to cold exposure. After cold exposure, the same pots were retained, but the lids were replaced with mesh secured with an elastic band to allow adult emergence. Pots were then returned to colony conditions, and survival was assessed on day 6 post-exposure, based on the number of individuals that successfully emerged as adults (Table 1).

#### 2.4.6. Adults

Adults were anaesthetised using a Flowbuddy benchtop CO₂ regulator connected to a standard fly pad. The mesh cage containing adults was placed directly onto the pad, allowing CO₂ to pass through and immobilise individuals without injury. Adults were then sexed under mild anaesthesia and transferred into 125 ml pots, with 25 individuals per pot. Each pot contained 12 or 13 individuals of each sex, resulting in a balanced sex ratio across the full sample (n = 100 total; 50 females, 50 males). Following CO₂ exposure, adults were allowed to recover for 1 hour at ambient temperature before being subjected to cold treatments. After cold exposure, pots were again held at ambient temperature for 1 hour to allow recovery. Lids were then replaced with mesh, and pots were returned to colony conditions. Survival was recorded 1 hour post-recovery and again after three days (Table 1). During this period, moist cotton wool soaked in 10% sucrose solution was placed on top of the mesh to provide continuous access to food.

### 2.5 Temperature treatments

#### 2.5.1 One hour exposure experiments

Each life stage was subjected to a series of acute (1 hour) temperature exposures ranging from −1°C to −10°C, in 1°C increments, without prior acclimation, as a previous study found no significant differences in cold tolerance with or without acclimation (McDermott et al. 2017). If survival was observed at −10°C, additional lower temperatures were tested to determine the lethal limit for that life stage. The lowest temperature that could be tested was −18°C, which represented the lower operational limit of the water bath. Therefore, the lower thermal limits reported for eggs should be interpreted with caution.

#### 2.5.2 Extended duration exposure experiments (6 and 24 hours)

Temperatures for the 6 hour and 24 hour exposures were selected based on life stage-specific survival patterns observed in the 1 hour trials. For each life stage, three or four representative temperatures were chosen to capture a range of survival outcomes. These included −1 °C as the upper end of the subzero range, a mid-range temperature that produced partial mortality (typically ∼30–60% survival), and one or two temperatures just above the point at which survival approached zero (∼0–1%; Table 2). If individuals survived exposure to −10°C in the 1 hour trials, they were subsequently tested at lower temperatures until no survival was detected. This approach allowed for the identification of life stage-specific cold tolerance thresholds. Therefore, the lower temperature ranges differed among life stages (accounted for statistically). The exception was the egg stage, where survival was still observed at −18°C. The system could not reliably maintain temperatures below this point, and at −18°C stability of the water bath was limited to one hour; therefore, for longer exposures the lowest consistently maintained temperature was −14°C. For pupae, four temperature treatments were used rather than three because complete mortality occurred at the lowest selected temperature during the 6 hour exposure. An additional temperature was therefore included to provide finer resolution immediately above this threshold and to more accurately characterise the pupal cold tolerance limits. Following cold exposure, specimens were allowed to recover for 1 hour at ambient room temperature (approximately 20±2°C) before being returned to colony conditions. Experiments were conducted in a randomised order across different times of the day under standard artificial laboratory lighting.

**Table 2.** Temperature exposures for each life stage of *Culicoides nubeculosus* across 1 hour, 6 hour, and 24 hour cold exposure trials. One-hour trials included a wide temperature range (down to the lower operational limit of −18°C) to determine acute cold tolerance. For the 6 and 24 hour trials, a subset of temperatures was selected based on low, moderate, and high survival observed in the 1-hour exposures. All tests included a control group.

| Life stage | 1 hour temperatures tested | 6 and 24 hour temperatures tested |
| --- | --- | --- |
| Eggs | Control, –1°C to –10°C (1°C increments); –11°C, –12°C, –14°C, –18°C | Control, –1°C, –11°C, –14°C |
| L1 larvae | Control, –1°C to –10°C (1°C increments) | Control, –1°C, –6°C, –10°C |
| L4 larvae | Control, –1°C to –10°C (1°C increments) | Control, –1°C, –6°C, –7°C |
| Pupae | Control, -1°C to -10°C (1°C increments); -11°C, -12°C, -13°C, -14°C | Control, -1°C, -8°C, -10°C, -14°C |
| Adults | Control, -1°C to -10°C (1°C increments) | Control, -1°C, -5°C, -10°C |

### 2.6 Wing Dissection and Measurement

Individuals that developed to the adult stage from each prior life stage were collected for wing measurements. When fewer than 20 adults survived to emergence, all were measured; when more than 20 survived, a random subset of 20 individuals were measured, aiming for ∼5 per replicate. If a replicate yielded fewer than 5 adults, all were included and additional adults were measured from other replicates where possible to approach a total of 20 individuals per life stage, which was achieved in approximately 40% of cases. In instances where this was not achievable due to low overall survival, all available individuals were measured, resulting in final sample sizes ranging from 1 to 20 individuals. The left wing of each adult was carefully dissected using fine forceps under a microscope and mounted on a microscope slide. Wing images were captured, and measurements were taken using the straight-line tool in ImageJ software (Schneider et al. 2012) from the arculus to the distal tip of vein M1. This linear distance was used as a consistent proxy for overall body size. If the left wing was damaged, the right wing was used instead. Wings that were damaged or incomplete were excluded from analysis.

### 2.7 Statistical analyses

All statistical analyses were conducted in R version 4.4.2 (R Core Team, 2024) to assess the effects of temperature and exposure duration on survival, hatching, development, emergence, and wing length across multiple life stages. Depending on the structure of the data and the research question, generalised linear models (GLMs) or mixed-effects models (generalised linear mixed models, GLMMs, or linear mixed-effects models, LMMs) were applied, including both additive and interaction terms where appropriate. Additive models were used where interaction terms were non-estimable because of missing data or zero emergence in certain treatment combinations (e.g. wing-length and adult-emergence models). Temperature and exposure duration (1, 6, or 24 hours) were treated as fixed effects in all models. In models assessing wing length, sex was included as an additional fixed factor. Random intercepts for replicate were incorporated specifically in time-series hatching models (hatching rate across days) and morphometric analyses (wing length) using mixedeffects models (GLMMs for binomial hatching responses and LMMs for Gaussian winglength data), as these data structures exhibited repeated measures or nested clustering within replicates. Survival and emergence models were analysed using GLMs without random effects, as responses were assessed at single time points within each experimental unit (Supplementary Table S2). Binary response variables (i.e. survival, hatching, emergence and adult development) were modelled using binomial or quasibinomial families, while continuous traits (wing length) were analysed using the Gaussian family. Overdispersion was evaluated in each model using the DHARMa package and, where necessary, addressed using quasibinomial models (e.g. in the across-life-stage survival model and the L1 survival model). In cases of complete or quasi-complete separation, Firth-adjusted bias reduction was applied using the brglmFit method from the brglm2 package. Temperature was generally treated as a categorical variable to reflect the experimental design and to include the control treatment. However, in the across-life-stage survival model and the across-life-stage wing length model (Supplementary Table S2), temperature was modelled as a continuous variable to capture the overall trend across the −1 to −10 °C gradient. In the across-life-stage survival model, the control group was excluded to maintain the integrity of the temperature gradient. Statistical significance of fixed effects and interactions was assessed using Type III analysis of deviance using the car package. For the across-life-stage survival model and the larval instar 1 (L1) survival model, both of which used quasibinomial families due to overdispersion, Type III F-tests were applied instead. Model comparison and simplification were based on analyses of deviance and, where applicable, likelihood-ratio tests. Tukeyadjusted post hoc comparisons were performed using the emmeans package. All analyses were conducted using the following R packages: car, emmeans, DHARMa, brglm2, lme4, multcomp, ggeffects, ggplot2, cowplot, patchwork, viridis, dplyr, tidyr, stringr, tibble, purrr, grid, and gtable.

## 3. Results

### 3.1 All life stages after 1 hour cold exposure

Survival following a 1 hour exposure to sub-zero temperatures (−1 to −10 °C) differed significantly among life stages, temperatures, and their interaction (Figure 2; Table 3). Model predictions indicated that egg survival remained consistent across temperatures, pupae and adults showed moderate declines, and larval stages particularly L4 declined most sharply (Figure 2). Observed survivability thresholds reflected these patterns. Eggs survived to at least −18 °C, which was the lowest temperature tested and the technical limit of the water bath, so their true lower lethal limit could not be determined. Pupae survived down to −14 °C, and adults and L1 larvae survived to −10 °C. L4 larvae were the least cold-tolerant and survived only to −7 °C. The overall decline in rate of survival with decreasing temperatures aligned with these patterns (Figure 2). Predicted slopes of survival (log odds °C⁻¹), derived from the fitted binomial model, were steepest for L4 larvae (−0.99), followed by L1 larvae (−0.45), pupae (−0.25), and adults (−0.15), and near zero for eggs (−0.002). At −10 °C, both observed and predicted egg survival were significantly higher than those of larvae and adults (all p < 0.001), and pupal survival was significantly higher than both larval stages (p < 0.001). L4 larvae had the lowest overall tolerance.

**Figure 2.**
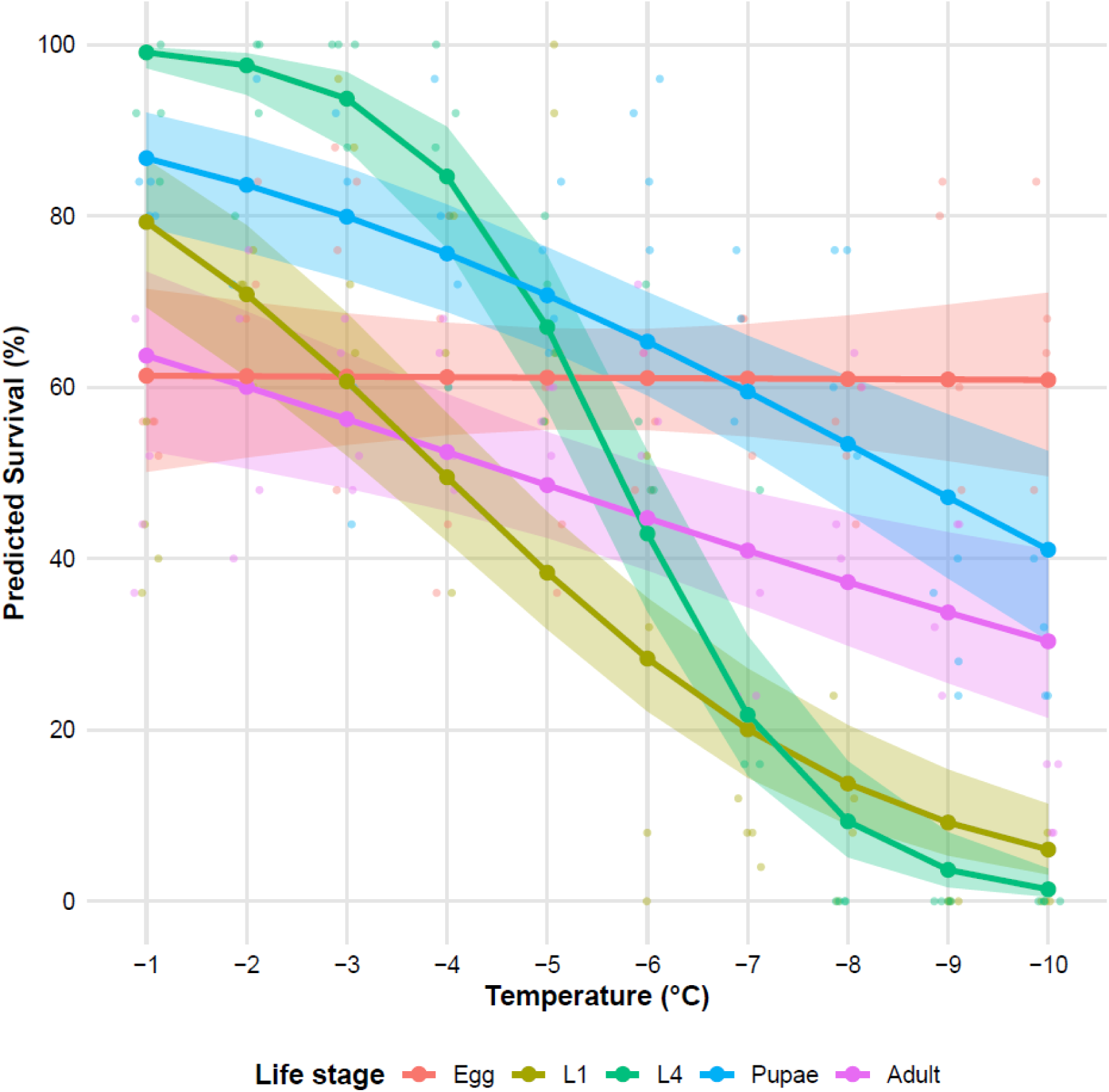
Predicted survival of *Culicoides nubeculosus* across developmental stages following a 1 hour exposure to sub-zero temperatures (−1°C to −10°C). Lines show estimated marginal means with 95% confidence intervals (shaded ribbons). Developmental stages include egg, larval instars (L1 and L4), pupae, and adults, color-coded as shown in the legend. Observed data points are overlaid as semi-transparent dots.

**Table 3.**
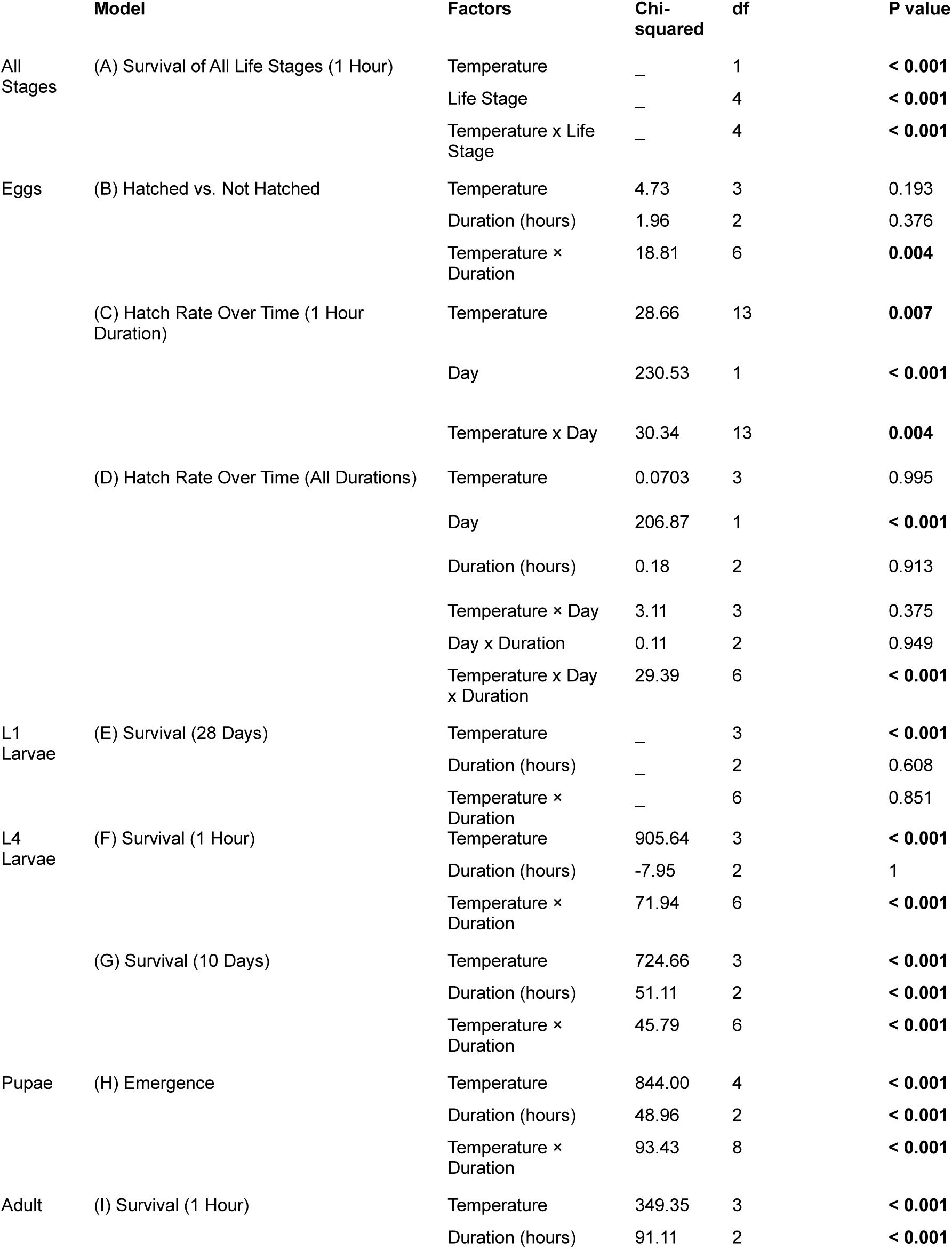
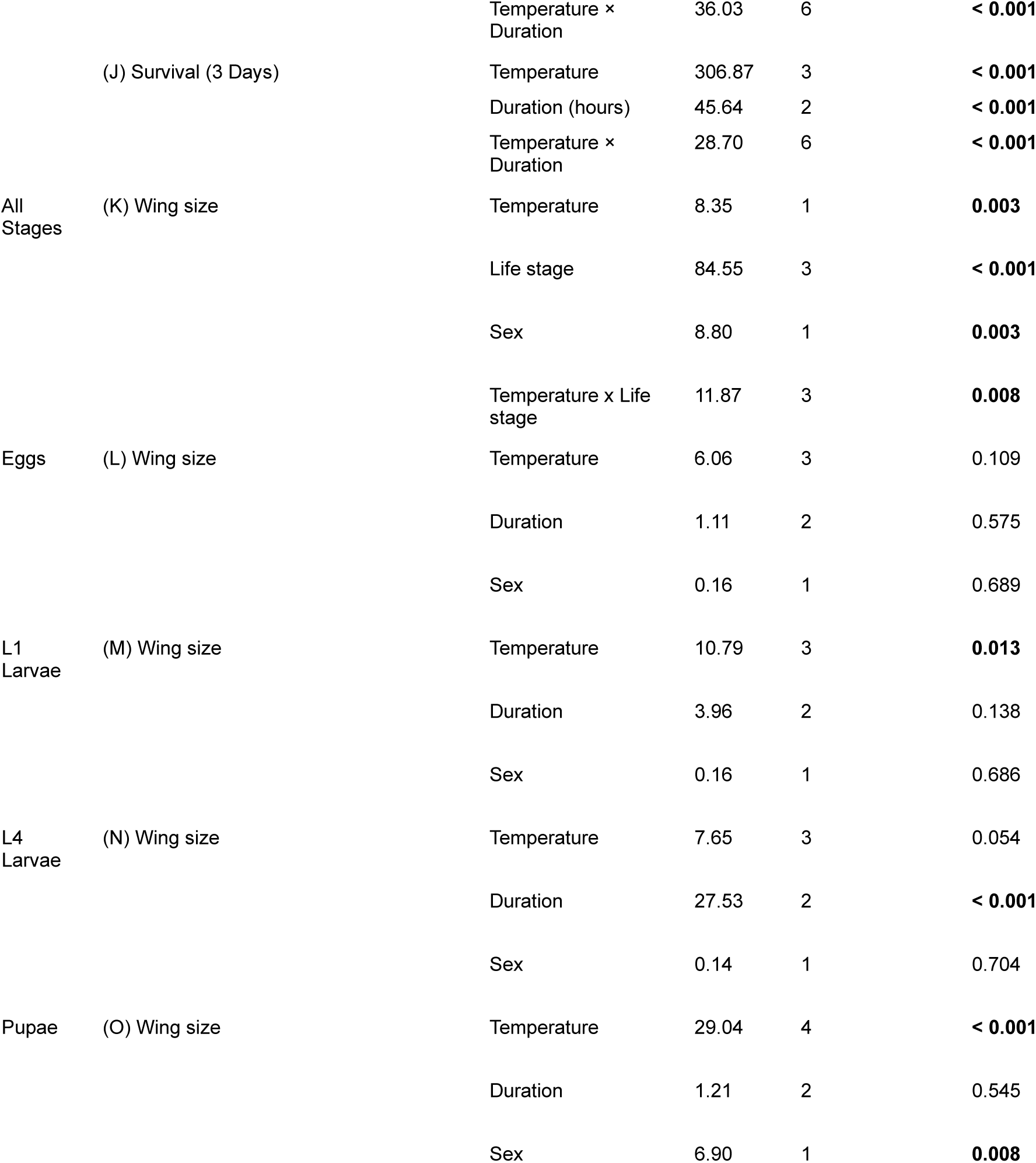
Analysis of deviance from generalised linear models (GLMs), generalised linear mixed models (GLMMs), and linear mixed models (LMMs) assessing the effects of temperature, exposure duration, and their interaction on survival and wing size in different life stages of *Culicoides nubeculosus* (Models A–O).

**Alt text:** Graph showing predicted survival of eggs, larvae, pupae and adults of *Culicoides nubeculosus* when exposed to temperatures of −1°C to −10°C for 1 hour.

### 3.2 Eggs

#### 3.2.1 Hatching success

Hatching success after 28 days was significantly influenced by an interaction between temperature and exposure duration (Table 3). Significant differences between temperature treatments were only observed after 24 h exposure, where hatching proportions were higher at −1 °C (p = 0.006) and −11 °C (p = 0.037) compared to control, while the difference at −14 °C was not significant (Figure 3). No significant temperature differences were observed for exposure times of 1 h or 6 h. There were no significant differences between the durations when temperature groups were pooled (Figure 3).

**Figure 3.**
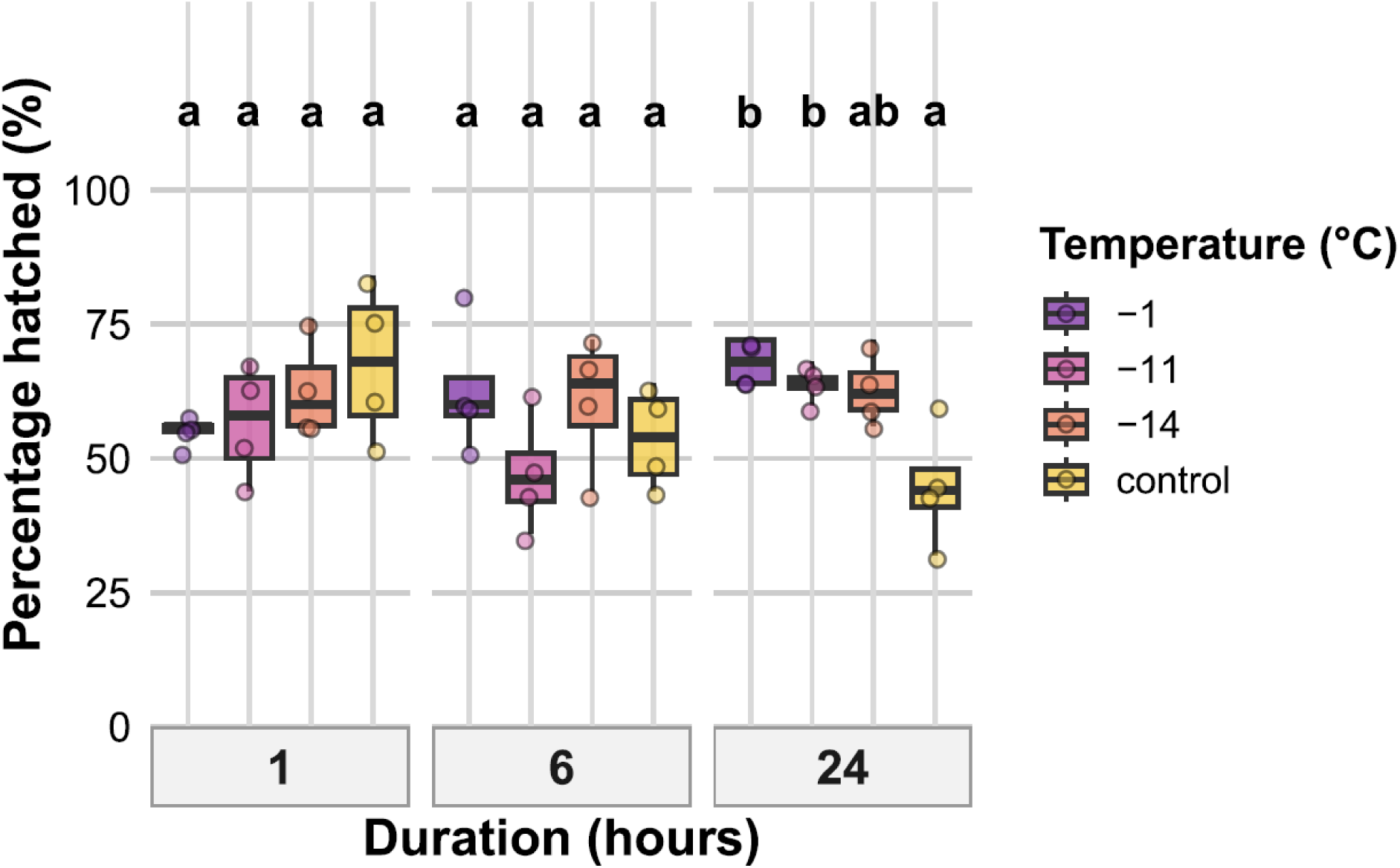
Effects of temperature and exposure duration on hatching success after 28 days on *Culicoides nubeculosus* cold exposed eggs. Boxplots show percentage of eggs that hatched across four temperature treatments (−1 °C, −11 °C, −14 °C, and control) and three exposure durations (1, 6, and 24 hours). Coloured points represent individual data. Boxplot colours correspond to temperature treatments as shown in the legend. Facet panels indicate different exposure durations. Lowercase letters above boxplots indicate statistically significant differences between temperature treatments within the same exposure duration. Treatments sharing the same letter within a duration are not significantly different (p > 0.05). The y-axis represents percentage values.

**Alt text:** Graph showing the egg hatch rate as a percentage for *Culicoides nubeculosus* eggs exposed to temperatures of −1 °C, −11 °C, −14 °C, and control for 1, 6 and 24 hour durations.

#### 3.2.3 Proportion of eggs hatched over time (1 hour exposure)

After 1 hour of cold exposure, hatching success over time was significantly influenced in interaction with temperature (Table 3). Eggs exposed to −18 °C and those in the control group (i.e., the lowest/highest temperature extremes) hatched faster, showing the steepest increases in hatch proportion over time. Slopes decreased progressively at −14 °C, −11 °C, and −1 °C, but all were significantly greater than zero (p < 0.05; Figure 4A), indicating that hatching continued to increase over time under all treatments.

**Figure 4.**
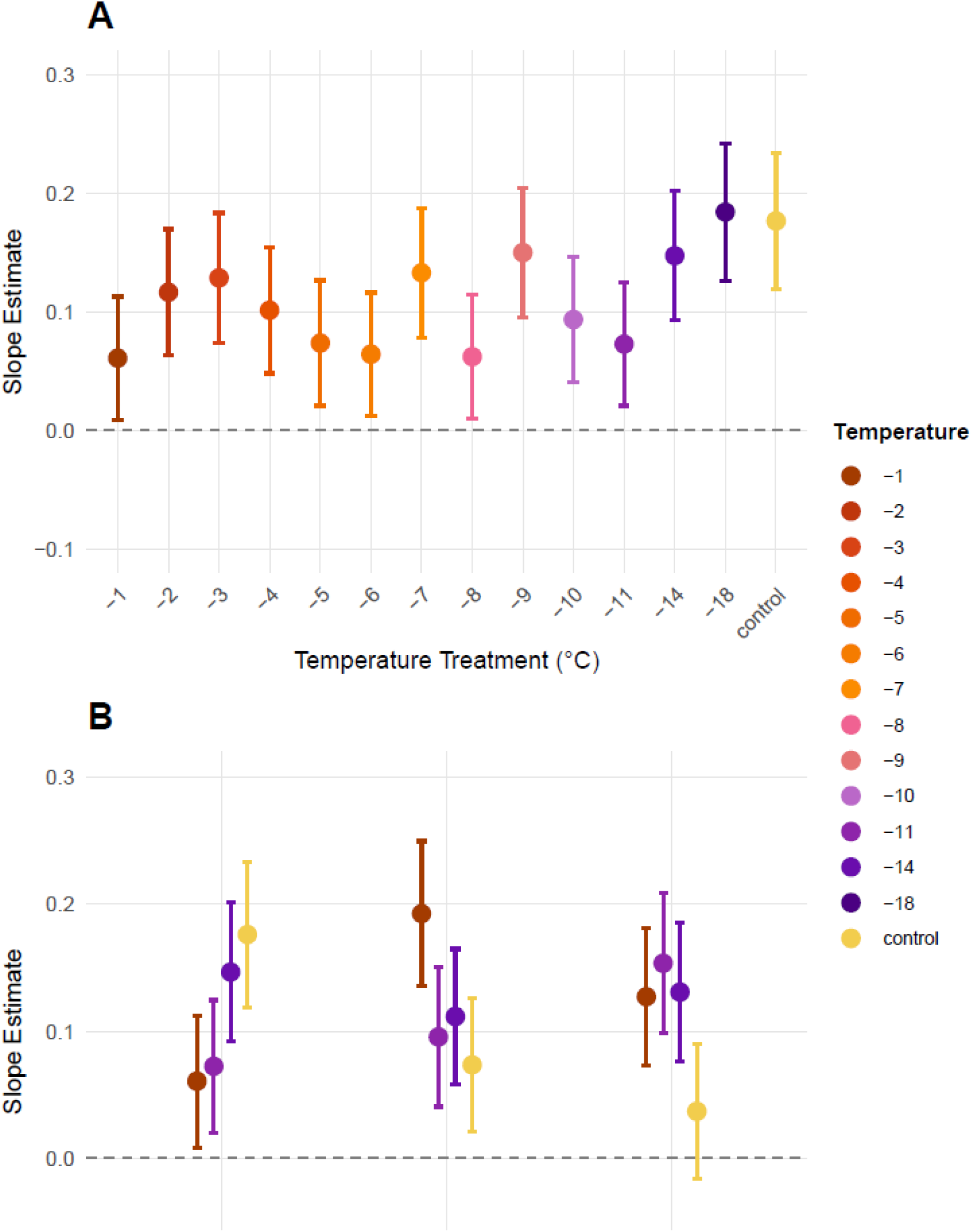
Temporal trends in egg hatch rate of *Culicoides nubeculosus* under cold exposure conditions. (A) Slope estimates (± 95% CI) describing the rate of increase in hatching over the full 14-day period following 1 hour exposure to a range of temperature treatments (−1°C to −18°C) and a control. (B) Slope estimates (± 95% CI) across 1 hour, 6 hour, and 24 hour exposure durations at selected temperatures (−1°C, −11°C, −14°C, and control).

#### 3.2.4 Proportion of eggs hatched over time across exposure durations

Across all exposure durations (1 h, 6 h, 24 h), hatch rate increased significantly over time, while temperature and duration did not have significant main effects. No significant two-way interactions were observed between temperature and time or between time and duration. However, a significant three-way interaction between temperature, day, and duration was found (Table 3). At −1°C, the slope for 6 hour exposure was significantly greater than at 1 h exposure (p = 0.039). No other durations showed significant pairwise differences within temperatures. In the control group, hatch rate increased significantly for 1 hour and 6 hours of exposure, but not for 24 hours of exposure, where the slope was not significantly different from zero (Figure 4B).

**Alt text:** Two graphical representations, one showing slope estimates for the increase in *Culicoides nubeculosus* egg hatching rate over 14 days following 1 hour cold exposure to temperatures ranging from −1 °C to-18 °C and the other showing slope estimates for 1, 6 and 24 hour exposure durations at −1°C, −11°C, −14°C, and control.

### 3.3 Larval instar 1 (L1)

#### 3.3.1 Survival

Survival of L1 larvae at 28 days after cold exposure was significantly affected by temperature (p < 0.001), but not by exposure duration or their interaction (Table 3). Across all exposure durations, survival was significantly lower at −6 °C and −10 °C compared to the control, while survival at −1 °C was not significantly different from the control (Figure 5).

**Figure 5.**
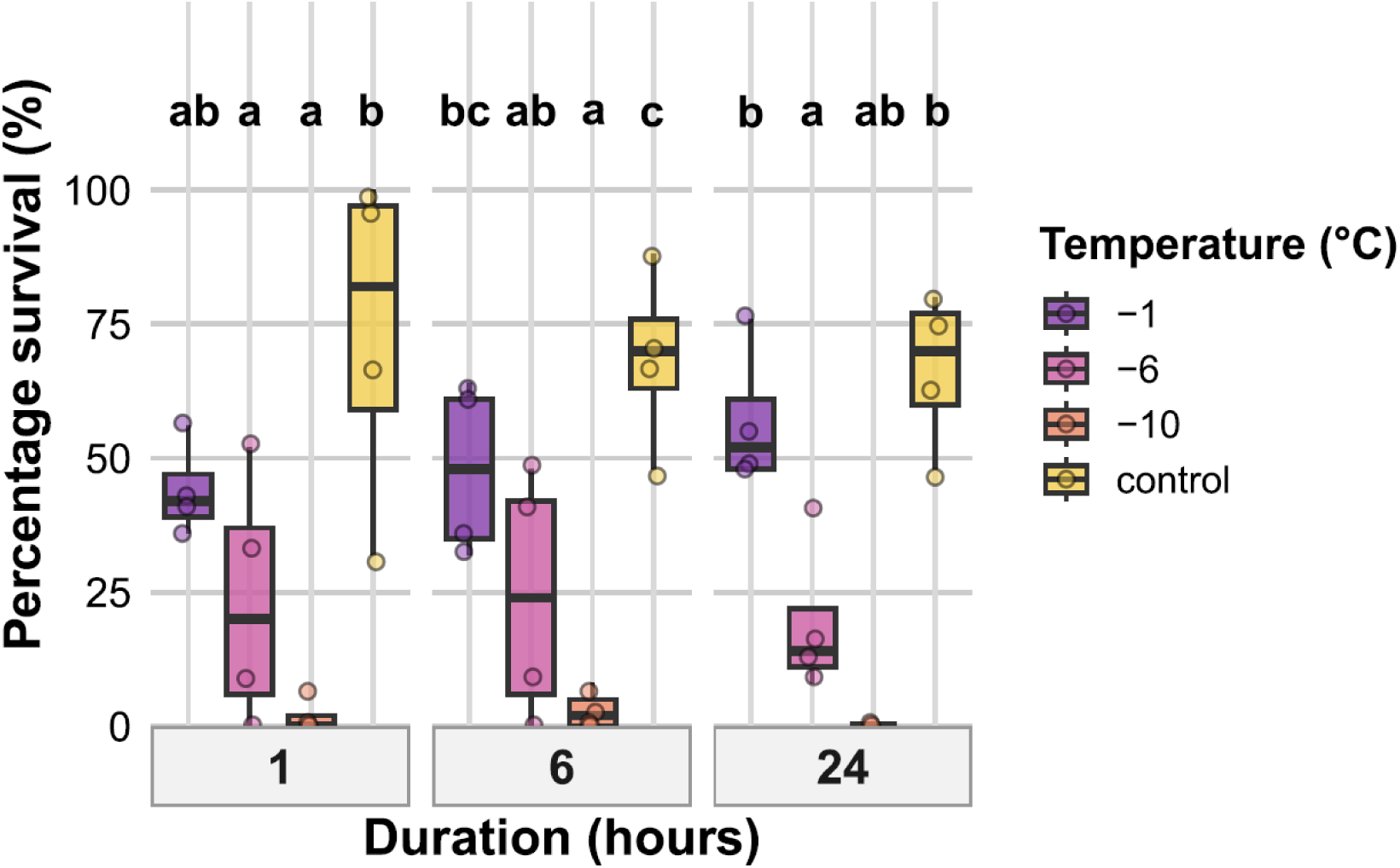
Effects of temperature and exposure duration on survival of *Culicoides nubeculosus* L1 larvae at 28 days post-exposure. Boxplots show percentage survival across four temperature treatments (−1 °C, −6 °C, −10 °C, and control) and three exposure durations (1, 6, and 24 hours). Coloured points represent individual data. Boxplot colours correspond to temperature treatments as indicated in the legend. Facet panels indicate different exposure durations. Lowercase letters above boxplots denote statistically significant differences between temperature treatments within each duration; treatments sharing the same letter are not significantly different (p > 0.05).

**Alt text:** Graph showing percentage survival of L1 larvae at 28 days after exposure to temperatures of −1 °C, −6 °C, −10 °C, and control for 1, 6 and 24 hour durations.

### 3.4 Larval instar 4 (L4)

#### 3.4.1 Survival after 1 hour

Initial survival of L4 larvae at 1 hour post-exposure was significantly affected by the interaction between temperature and exposure duration (Table 3). After 1 hour cold exposure, survival at −7 °C was significantly lower than all other treatments (p < 0.001), while survival at −6 °C was significantly reduced compared to both −1 °C and control (p = 0.004). For 6 hours of exposure, there was no survival at −7 °C, and this was significantly lower than survival at −6 °C, −1 °C, and the control (p < 0.001). Survival at −6 °C was also significantly lower than at both −1 °C and the control (both p = 0.042). For 24 hours of exposure, survival at −6 °C was significantly lower than at −1 °C and the control (both p = 0.002), while survival at −7 °C again showed no survivors and was significantly lower than both −1 °C and the control (p < 0.001). Survival at −6 °C and −7 °C also consistently differed significantly from one another (p < 0.001). There were no significant differences between the durations when temperature groups were pooled (Figure 6A).

**Figure 6.**
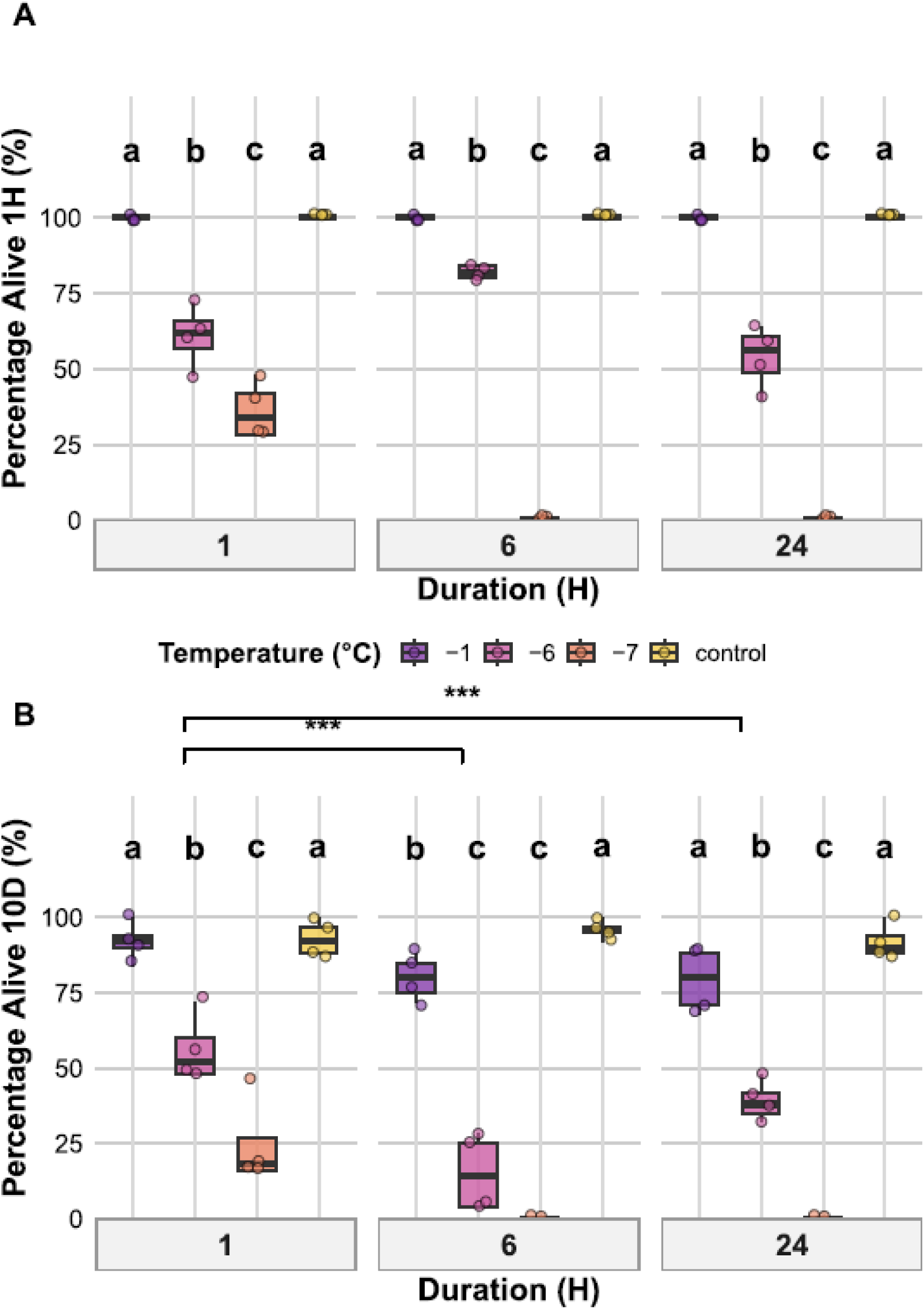
Effects of temperature and exposure duration on L4 larval *Culicoides nubeculosus* survival. Boxplots show (A) percentage of larvae alive after 1 hour post-exposure, (B) percentage alive after 10 days post-exposure across four temperature treatments (−1 °C, −6 °C, −7 °C, and control) and three exposure durations (1, 6, and 24 hours). Coloured points represent individual data. Boxplot colours correspond to temperature treatments as shown in the legend. Facet panels indicate different exposure durations. Lowercase letters above boxplots indicate statistically significant differences between temperature treatments within the same exposure duration. Treatments sharing the same letter within a duration are not significantly different (p > 0.05). The y-axis represents percentage values. Brackets with asterisks indicate significant differences between exposure durations when temperature treatments are pooled (control excluded): * p < 0.05, ** p < 0.01, *** p < 0.001.

#### 3.4.2 Survival after 10 days

At 10 days post-exposure, survival outcomes were significantly influenced by an interaction between temperature and duration (Table 3). For 1 hour exposure, survival at −7 °C was significantly lower than at all other temperatures (p < 0.001), and survival at −6 °C was also significantly lower than both −1 °C (*p* < 0.001) and the control (*p* < 0.001). For 6 hours of exposure, survival at −7 °C dropped to zero, and survival at −6 °C was again significantly lower than both −1 °C and the control (both < 0.001). For 24 hours of exposure, survival remained high in the −1 °C and control groups, while no individuals survived under −7 °C and survival was reduced at −6 °C, which was significantly lower than both −1 °C (*p* < 0.001) and control (*p* < 0.001). When temperature treatments were pooled (excluding control), survival after 10 days following 1 hour exposure was significantly higher than for both 6 and 24 hours of exposure (p < 0.001), while the difference between 6 and 24 hours of exposure was not significant (Figure 6B).

**Alt text:** Graphs showing the percentage of *Culicoides nubeculosus* L4 larvae surviving at 1 hour and 10 days post cold exposure to temperatures of −1 °C, −6 °C, −7 °C, and control for durations of 1, 6 and 24 hours.

### 3.5 Pupae

#### 3.5.1 Pupal Emergence Success

Adult emergence from pupae was assessed following exposure to temperature treatments for 1, 6, or 24 hours and measured after 6 days. Emergence was significantly affected by temperature, duration, and their interaction (Table 3). For 1 hour exposure, emergence under−1 °C and control conditions was significantly higher than −10 °C and −14 °C (*p* < 0.001).

Emergence under −8 °C was also significantly higher than under −10 °C and −14 °C (*p* < 0.001) but did not differ significantly from −1 °C or control. After 6 hours of exposure, emergence was highest at –1 °C and in the control, which did not differ significantly from one another. Both treatments showed significantly greater emergence than –8 °C, –10 °C, and – 14 °C (p < 0.001). Emergence at –8 °C was significantly higher than at –10 °C (p = 0.024) and –14 °C (p = 0.040), while emergence at –10 °C and –14 °C did not differ significantly. For 24hour exposures, emergence at –1 °C and in the control remained high and did not differ significantly. Emergence at –8 °C was significantly lower than both –1 °C and the control (p < 0.001), and emergence was absent at both −10 °C and −14 °C (Figure 7). When pooled across temperature treatments (excluding the control), pupal emergence was significantly reduced following 6 and 24 hour exposures compared to 1 hour (*p* < 0.001), with no significant difference observed between 6 and 24 hours.

**Figure 7.**
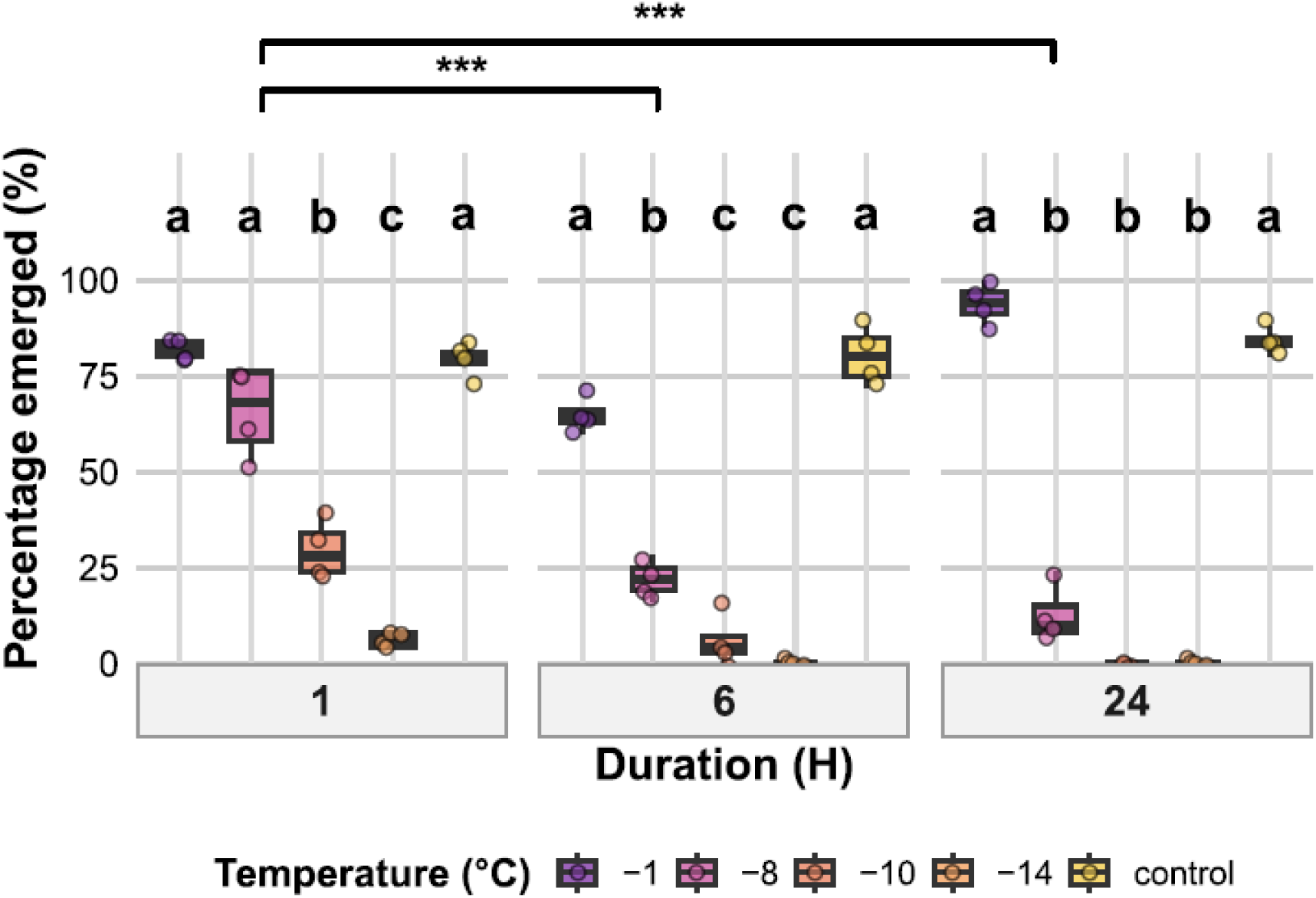
Effects of temperature and exposure duration on *Culicoides nubeculosus* adult emergence from pupae. Boxplots show the percentage of pupae that successfully emerged across five temperature treatments (control, −1 °C, −8 °C, −10 °C, and −14 °C) and three exposure durations (1, 6, and 24 hours). Coloured points represent individual data. Boxplot colours correspond to temperature treatments as shown in the legend. Facet panels indicate different exposure durations. Lowercase letters above boxplots indicate statistically significant differences between temperature treatments within the same exposure duration, based on post hoc Tukey-adjusted pairwise comparisons. Treatments sharing the same letter within a duration are not significantly different (p > 0.05). The y-axis represents percentage values. Brackets with asterisks denote significant differences between exposure durations (pooled across temperature treatments, excluding control): * p < 0.05, ** p < 0.01, *** p < 0.001.

**Alt text:** Graph showing the percentage of *Culicoides nubeculosus* pupae that emerged as adults after exposure to temperatures of −1 °C, −8 °C, −10 °C, −14 °C, and control for durations of 1, 6 and 24 hours.

### 3.6 Adults

#### 3.6.1 Adult survival after 1 hour

Adult survival at 1 hour post-exposure was significantly affected by temperature, duration, and their interaction (Table 3). Within each duration, survival varied significantly among temperatures (Figure 8A). For both 1 hour and 6 hour exposures, survival at −10 °C was significantly lower than all other temperatures (p < 0.001), while −1 °C, −5 °C, and control did not differ significantly. For 24 hours of exposure, survival at −10 °C was significantly lower than all other treatments (p < 0.001), and survival at −1 °C was significantly lower than at −5 °C (p < 0.001) and control (p < 0.001), while −5 °C and control did not differ. When temperature treatments were pooled (excluding the control), survival after 24 hours was significantly lower than after both 1 hour and 6 hours (p < 0.001), while survival after 1 and 6 hours did not differ.

**Figure 8.**
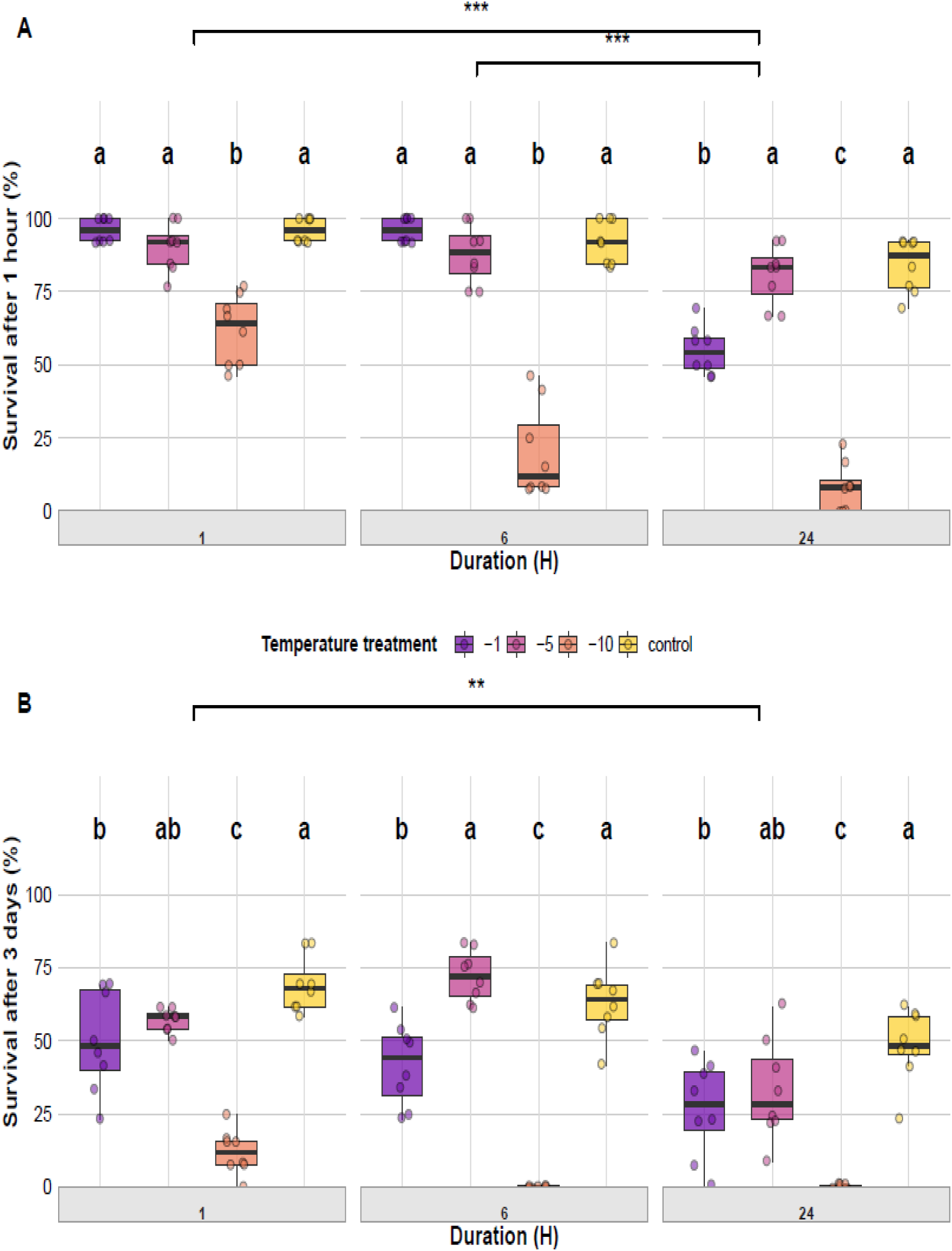
Effects of temperature and exposure duration on adult survival of *Culicoides nubeculosus*. Boxplots show the percentage of adults alive after either 1 hour (A) or 3 days (B) post-exposure across four temperature treatments (control, −1 °C, −5 °C, and −10 °C) and three exposure durations (1, 6, and 24 hours). Coloured points represent individual data, and box colours correspond to temperature treatments as indicated in the legend. Facet panels denote different exposure durations. Lowercase letters above boxplots indicate statistically significant differences among temperature treatments within the same duration; treatments sharing a letter are not significantly different (p > 0.05). Brackets with asterisks show significant differences between exposure durations when temperature treatments were pooled (control excluded): * p < 0.05, ** p < 0.01, *** p < 0.001.

#### 3.6.2 Adult survival after 3 days

Adult survival at three days post-exposure was significantly influenced by temperature, duration, and their interaction (Table 3). Survival declined sharply with decreasing temperature at all exposure durations, with the lowest survival consistently observed at −10

°C (Figure 8B). For 1 hour exposure, survival at −10 °C was significantly lower than at −1 °C (p < 0.001), −5 °C (p < 0.001), and the control (p < 0.001), and survival at −1 °C was also lower than at the control (p = 0.036). For 6 hours of exposure, survival at −10 °C was again significantly lower than at −1 °C (p = 0.003), −5 °C (p < 0.001), and the control (p < 0.001); survival at −1 °C was lower than at −5 °C (p < 0.001) and the control (p = 0.018). After 24 hours of exposure, no individuals survived at −10 °C, with significantly higher survival at −1 °C (p = 0.014), −5 °C (p = 0.008), and the control (p = 0.002). Survival at −1 °C was also lower than the control (p = 0.014). When temperature treatments were pooled, duration also affected survival. Survival after 24 hours of exposure was significantly lower than after 1 hour (p = 0.001), while survival after 6 hours did not differ from either 1 hour or 24 hours.

**Alt text:** Graphs showing the percentage of *Culicoides nubeculosus* adults surviving at 1 hour and 3 days post cold exposure to temperatures of −1 °C, −5 °C, −10 °C, and control for durations of 1, 6 and 24 hours.

### 3.7 Wing Size

#### 3.7.1 Adult wing size after cold exposure of immature life stages (1 hour exposure)

The interaction between life stage at time of exposure and temperature had a significant effect on adult wing length (Table 3). This indicates that the effect of temperature on wing size differed among life stages. Only L4 larvae showed a clear temperature-dependent change in adult wing length, with wing length increasing as exposure temperature decreased (−21.6 µm per °C; p < 0.001, ≈1.5–2% increase per degree of cooling). No such temperature effect was detected for cold-exposed eggs, L1 larvae or pupae. Wing length also differed significantly among life stages (Table 3). Adults from immatures that were cold exposed at the egg stage had the smallest wings (1345 ± 125 µm), whereas those from cold-exposed L1 larvae were the largest (1481 ± 136 µm), with L4 larvae (1384 ± 126 µm) and pupae (1459 ± 127 µm) intermediate. Pairwise post-hoc comparisons showed that adults from eggs had significantly smaller wings than those from all other life stages (Egg vs L1: p < 0.001; Egg vs L4: p = 0.005; Egg vs Pupae: p < 0.001). Wings of adults from L1 larvae were significantly larger than those from L4 larvae (p < 0.001), while pupal adults were intermediate, with wings significantly larger than those from L4 larvae (p = 0.009) but not significantly different from those from L1 larvae (p = 0.31). Females had slightly longer wings than males (1434 vs. 1401 µm; ≈2–3% larger) across all treatment groups (Figure 9).

**Figure 9.**
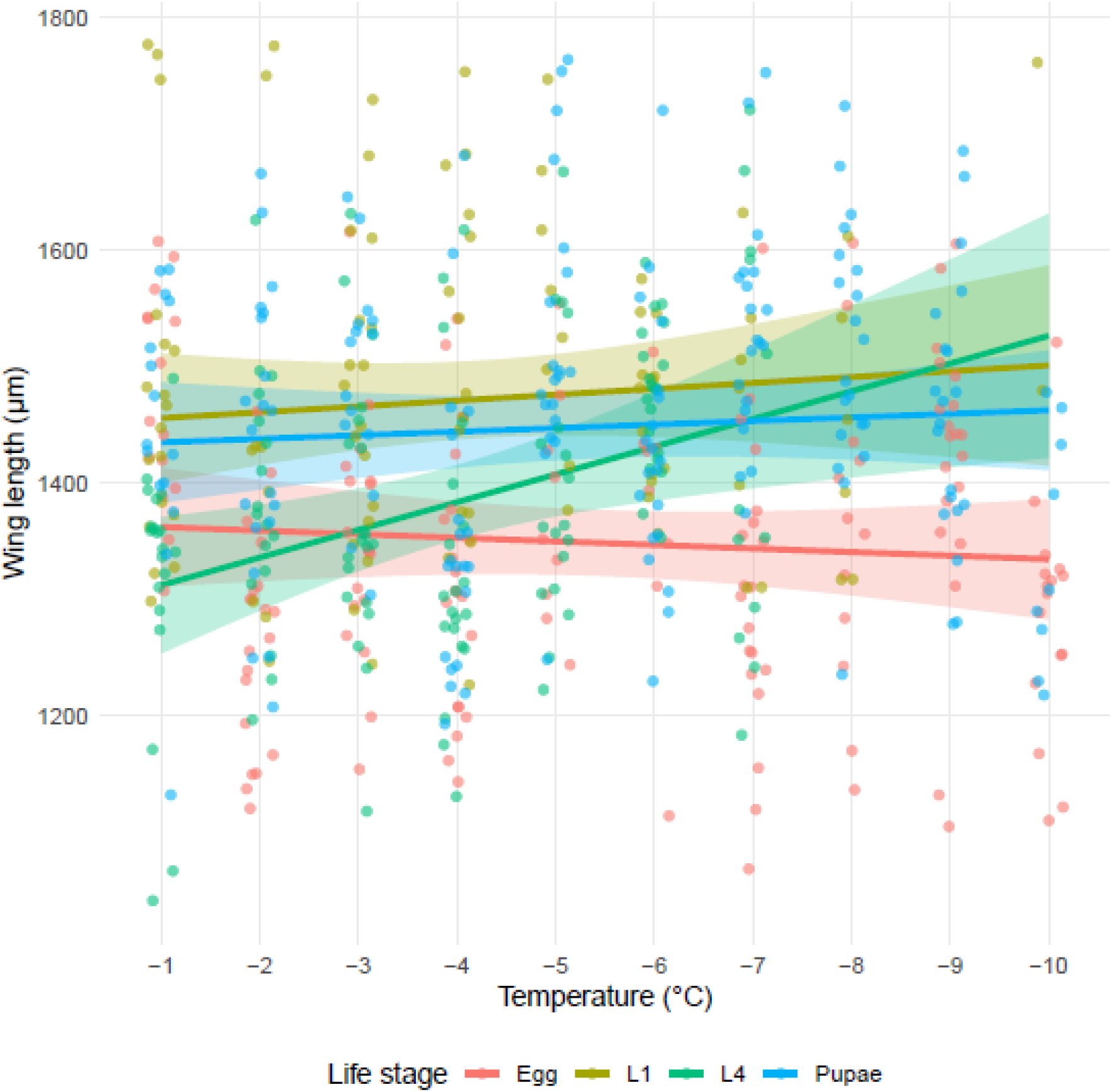
Wing length of *Culicoides nubeculosus* adults following a 1 hour exposure to subzero temperatures (−1 °C to −10 °C) during different developmental stages. Lines represent fitted regression models with 95% confidence intervals (shaded ribbons). Developmental stages exposed include egg, larval instars (L1 and L4), and pupae, color-coded as shown in the legend. Observed wing length measurements are plotted as semi-transparent points.

**Alt text:** Graph showing the wing length of adult *Culicoides nubeculosus* after being exposed as eggs, larvae or pupae to temperatures ranging from −1 °C to −10 °C.

#### 3.7.2 Wing size after egg cold exposure

Wing length of adults exposed at the egg stage was not significantly affected by temperature, exposure duration, or sex (Table 3) indicating that cold exposure during the egg stage did not measurably influence adult wing size. Mean wing lengths were similar across all treatments (Figure 10A).

**Figure 10.**
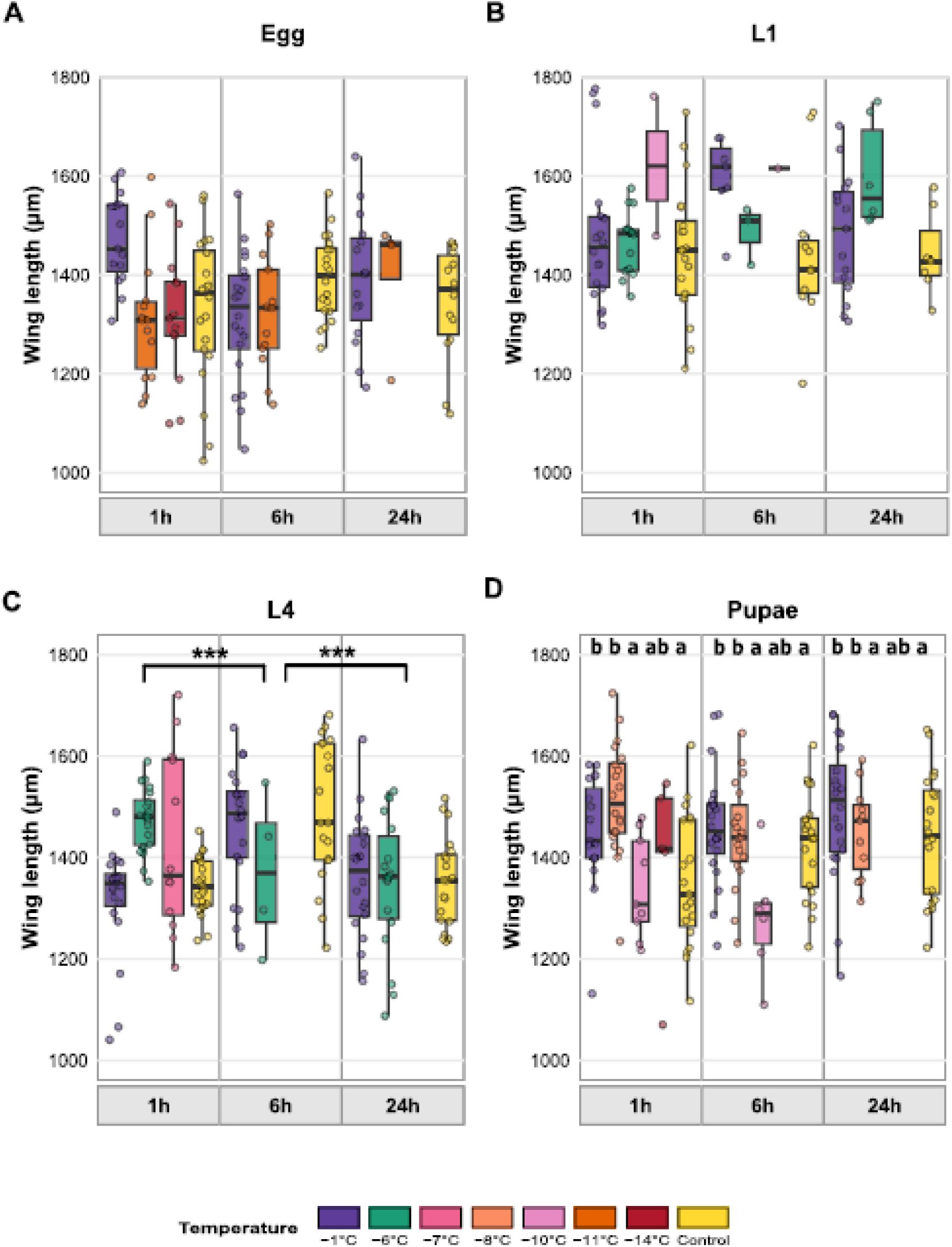
Effects of temperature and exposure duration on adult wing length. Boxplots show wing length (µm) of adult *Culicoides nubeculosus* after cold exposure at the (A) egg, (B) L1 larval stage, (C) L4 larval stage, and (D) pupal stages across differing temperature treatments (control, −1 °C, −6 °C, −7 °C, −8 °C, −10 °C, −11 °C and −14 °C) and three exposure durations (1, 6, and 24 hours). Coloured points represent individual data. Boxplot colours correspond to temperature treatments as shown in the legend. Facet panels indicate different exposure durations. For L4 larvae (C), bracketed asterisks indicate the main effect of exposure duration, averaged across cold-temperature treatments (control excluded). For pupae (D), lowercase letters indicate the main effect of temperature: temperatures that do not share a letter differ significantly when averaged across durations, and the same groupings are shown in each duration panel. No significant overall temperature effects were detected for eggs, L1 or L4 larvae, so letters are not shown. Significance: *p < 0.05, **p < 0.01, ***p < 0.001.

#### 3.7.3 Wing size after L1 larval cold exposure

Wing length of adults exposed at the L1 larval stage was significantly affected by temperature, but not by exposure duration or sex. Mean wing lengths (estimated marginal means) ranged from 1439 ± 36 µm (control) to 1615 ± 78 µm (−10 °C), with no significant differences among treatments (Figure 10B).

#### 3.7.4 Wing size after L4 larval cold exposure

Wing length of adults exposed at the L4 stage was significantly affected by both temperature and exposure duration (Table 3). Mean wing lengths ranged from 1380 ± 20 µm (−1 °C) to 1459 ± 40 µm (−7 °C) across temperature treatments, and from 1368 ± 23 µm (24 h) to 1490 ± 26 µm (6 h) across exposure durations. Sex had no significant effect on wing length (Table 3). Wings were longest following 6-hour exposures, being significantly longer than at 1 hour (p < 0.001) and 24 hours (p < 0.001; Figure 10C).

#### 3.7.5 Wing size after pupal cold exposure

Wing length of adults emerging from cold exposed pupae was significantly affected by temperature and sex, but not by exposure duration (Table 3, Figure 10D). Mean wing lengths ranged from 1317 ± 33 µm (−10 °C) to 1471 ± 17 µm (−8 °C) across temperature treatments, with individuals at −1 °C and −8 °C having significantly longer wings than those at −10 °C (p = 0.001 and p < 0.001, respectively) and longer wings than controls (p = 0.037 and p = 0.010, respectively). Females had significantly larger wings (1434 ± 16 µm) than males (1385 ± 19 µm; p = 0.012).

**Alt text:** Four graphs showing wing length of adult *Culicoides nubeculosus* after cold exposure as eggs, larvae and pupae to temperatures of −1 °C, −6 °C, −7 °C, −8 °C, −10 °C, −11 °C, −14 °C, and control for durations of 1, 6 and 24 hours.

### 3.8 Developmental outcomes

Cold exposure during the egg, L1, and L4 stages significantly influenced the proportion of individuals that completed development to adulthood, with significant temperature × duration interactions in all three stages (Table S2). Across stages, adult emergence decreased at lower temperatures and longer exposure durations. Full analyses and figures are provided in the Supplementary Material (Section S3: Developmental Outcomes; Table S2; Figures S1– S2).

## Discussion

This study is the first to assess cold tolerance in a Palaearctic species of *Culicoides.* Across all life stages, *C. nubeculosus* displayed substantial cold tolerance. Eggs were consistently the most tolerant, followed by pupae, while larvae and adults showed lower tolerance but were still able to survive temperatures below those typically experienced in their natural environment. A similar life-stage pattern has been observed in *C. sonorensis*, where eggs tend to be the most cold tolerant and larvae are more sensitive (McDermott et al. 2017). The assessment of cold tolerance in *C. sonorensis* was based on acute 1 hour exposures of three immature stages (eggs, L4 larvae and pupae), whereas the present study examined additional life stages (adults and L1 larvae) and exposure durations (6 hours and 24 hours), providing a broader view of how cold tolerance varies throughout the *Culicoides* life cycle. Survival outcomes were strongly influenced by both temperature and duration of exposure, with colder conditions and longer exposures increasing mortality across all stages, although thresholds differed among them. These findings suggest that overwintering potential is not restricted to a single developmental stage and that cold snaps alone are unlikely to eliminate populations. This broad tolerance also has implications for geographical range expansion, as the capacity of multiple life stages to survive below typical thermal limits suggests that *C. nubeculosus* and other temperate *Culicoides* species may persist in cooler or more variable environments than currently recognised.

The cold tolerance observed across developmental stages of *C. nubeculosus* is likely supported by a range of physiological and structural traits. The ability of *Culicoides* eggs to survive cold exposure may be underpinned by mechanisms linked to desiccation tolerance in insects (Terhzaz et al. 2015). Since both freezing and desiccation cause cellular dehydration and membrane stress, physiological responses such as membrane stabilisation, cryoprotectant accumulation, and water regulation may contribute to cold tolerance in insects, including biting midges (Sinclair et al. 2013, McDermott and Mullens 2014, Terhzaz et al. 2015). Structural features including chorion tanning and small egg size may further reduce the likelihood of ice formation and mechanical damage (Cribb and Chitra 1998, McDermott et al. 2017). In *C. sonorensis* and related species, pupae gain protection from their hardened cuticle, which can limit water loss and reduce cold-induced damage (Gołębiowski et al. 2011). In contrast, larval stages of *Culicoides* and other Diptera often lack protective barriers and depend on substrate buffering, which leaves them more vulnerable to cold stress. Together, these mechanistic patterns emphasise that overwintering capacity depends not only on environmental conditions but also on stage-specific protective traits. In adult insects such as *Drosophila* and *C*. *sonorensis*, physiological mechanisms such as rapid cold-hardening can enhance short-term tolerance to subzero temperatures, allowing survival following brief preexposure to mild chilling (Nunamaker 1993). This rapid acclimation response likely involves membrane stabilisation, ion regulation, and the induction of heat-shock proteins that help maintain cellular function during thermal stress (Sinclair et al. 2013). Despite these protective responses, cold injury can still occur when temperatures fall below physiological limits, leading to neuromuscular impairment and increased levels of delayed mortality.

Cold exposure produced important sublethal consequences that extended beyond immediate survival, reflecting stress-induced physiological disruptions across stages. Eggs generally hatched successfully, even more so after extreme exposures, but reduced adult emergence revealed hidden fitness costs, a pattern also reported in other insects (Liao et al. 2021, Lü et al. 2025). Sublethal effects following cold exposure have also been noted in *C. sonorensis*, where egg and larvae surviving initial chilling later exhibit elevated delayed mortality (McDermott et al. 2017). In this study, at the lowest tested temperatures, hatching was accelerated following extreme cold, suggesting that cold stress induces a developmental stress response that promotes emergence once favourable conditions return. Although this has not been directly demonstrated in *Culicoides* prior to this study, similar environmentally cued plasticity in hatching timing has been reported in other dipterans. In mosquitoes, thermal and environmental cues can influence both the timing and proportion of hatching, allowing eggs to delay emergence under unfavourable conditions and resume development once conditions improve (Byttebier et al. 2014, Ebrahimi et al. 2014, Ottih and Tripet 2024). This suggests that the accelerated hatching observed in *Culicoides* eggs may reflect a similar adaptive mechanism promoting rapid emergence when conditions are favourable after cold stress.

Larvae also displayed delayed effects, with colder and longer exposures reducing successful pupation and adult emergence despite initial survival. Similar patterns of initial survival followed by later mortality have been documented in *C. sonorensis* larvae exposed to subzero temperatures, where individuals frequently died during subsequent development despite appearing viable immediately after chilling (McDermott et al. 2017). Observable stress responses were noted in L4 individuals during this study, which occasionally exhibited tremors and impaired movement following cold exposure, indicating chill coma-like effects — a transient neuromuscular dysfunction induced by low temperatures (MacMillan and Sinclair 2010, Overgaard and Macmillan 2017). Comparable effects have been reported in *Aedes aegypti* (Montini et al. 2021) and *Drosophila melanogaster* (Koštál et al. 2019), where larval cold stress delayed development and reduced adult fecundity. In our study, pupae tolerated short-term cold exposure with little immediate mortality, but prolonged exposure led to incomplete or failed metamorphosis, with fewer individuals successfully emerging as adults. Sublethal pupal injury and reduced emergence after cold exposure have similarly been observed in *C. sonorensis* pupae under short-term freezing stress (McDermott et al. 2017). These findings highlight the importance of assessing cold tolerance beyond immediate survival metrics, and it is possible that cross-tolerances may amplify the ability of insect vectors to withstand multiple stressors. Future research should explore whether these effects persist across generations, providing insights into the long-term adaptive potential of insect populations. It is worth nothing that differences in temperature thresholds for adult activity varied across the year in populations of field-caught *Culicoides* studied in the south of England, suggestive of population-level adaption to cold as temperatures drop in the autumn (Tugwell et al. 2021).

In this study, adults were the most visibly affected by cold stress, displaying tremors, impaired movement, and delayed mortality consistent with chill coma recovery. In several instances, individuals that initially recovered failed to survive the subsequent monitoring period, suggesting delayed physiological damage. The ability of some adults to survive brief exposure to −10 °C without prior acclimation suggests an inherent, if limited, resilience to acute cold shocks, though survival declined sharply with prolonged exposure. Moderate survival after 24 hours of exposure at –5 °C further suggests that short bouts of cold are tolerated better than prolonged exposure. This pattern reflects known physiological distinctions between acute and chronic chilling in chill-susceptible insects. The present results provide the first experimental data on how adult *Culicoides* respond to both acute and prolonged cold exposure. It is likely that the mechanisms of cold injury observed in other insects, are also applicable to *Culicoides.* While brief exposures may trigger reversible chill coma, extended cold exposure may progressively disrupt ion and water homeostasis across epithelial barriers such as the gut and Malpighian tubules. This can lead to cellular depolarisation, increased haemolymph potassium concentrations, and ultimately chilling injury and mortality (Macmillan et al. 2014, Overgaard and Macmillan 2017).

Although cold effects on wing size have not been examined previously in *Culicoides*, studies show that wing morphometrics vary among species and provide useful biological information (Slama et al. 2021, Hadj-Henni et al. 2023b, Hadj-Henni et al. 2023a). In this study, differences in wing size were generally small, but a temperature-related response was apparent in L4 larvae, suggesting that sensitivity to cold exposure may vary across life stages. Such stagespecific plasticity has been observed in other dipterans, where developmental temperature influences adult wing morphology. In mosquitoes, wing size is a reliable proxy for body size, and *Aedes albopictus* populations from cooler climates have larger wings, with larger size suggested to confer advantages such as increased energy reserves and improved cold tolerance (Sherpa et al. 2022). Similarly, in *Drosophila*, lower developmental temperatures consistently produce larger wings (Machida et al. 2022). These parallels suggest that the larger wings we observed in *C. nubeculosus* following cold exposure at the L4 stage may reflect a general ectothermic response to low developmental temperatures. Wing size may also influence survival, dispersal, and vectorial capacity, with larger wings potentially enhancing flight efficiency, longevity, and mating success, traits that can increase the likelihood of pathogen transmission (Alto et al. 2008, Juliano et al. 2014). In *Aedes aegypti*, larger females have been shown to live longer and exhibit higher infection rates, extending the potential transmission window for arboviruses (Juliano et al. 2014), although larger individuals can, in some cases, also exhibit stronger immune responses against the infecting pathogen. However, relationships between size, fecundity, and transmission are often nonlinear; larger individuals may face increased metabolic demands or delayed maturation, limiting reproductive output despite enhanced survival (Sherpa et al. 2022). In *Culicoides*, such trade-offs could determine whether cold-induced increases in wing size translate into a genuine fitness advantage or simply reflect developmental plasticity under thermal stress.

While laboratory assays provide valuable insights, they may either underestimate or overestimate survival in nature because they lack the buffering effects of natural substrates and the ecological and microbial complexity of real-world environments, including predator– prey interactions, competition, and diverse microbial communities (Kameke et al. 2021, Neupane et al. 2025). Larval performance under cold stress was poorer than that of other life stages, likely due to the limited thermal buffering provided by the polyester batting and small volume of rearing fluid used in this study (Devlin et al. 2022). Unlike natural substrates such as dung, soil, or compost, which retain heat and moisture and can generate additional microbial warmth (Lühken et al. 2014, Steinke et al. 2014, Erram et al. 2019, Chen et al. 2024, Hochstrasser et al. 2024), the experimental substrate provided minimal insulation. Previous studies have demonstrated that larvae can be found at varying depths in natural substrate, and it has been proposed that larvae can move vertically to regulate their exposure to temperature and moisture (Mullens and Rodriguez 1992, Blackwell and King 1997, Uslu and Dik 2006). Smaller L1 larvae may have gained greater protection by being more fully surrounded by the substrate or burrowing proportionally deeper, providing improved insulation and potentially higher survival than the larger, more exposed L4 larvae. Pupae similarly lacked insulation, as they were placed on damp filter paper rather than within protected microhabitats found in the field (Bishop and McKenzie 1994, Stokes et al. 2022) where substrates can provide greater thermal stability and support higher overwintering success (Van Den Eynde et al. 2021, Stokes et al. 2022). In addition, individuals in this study were returned to optimal colony conditions following cold exposure, which may have promoted recovery and overestimated post-exposure survival compared with field conditions, although the rapid transition from cold to warm temperatures may conversely have induced thermal shock and reduced survival, and indeed the exposure temperatures used are likely much lower than those experienced in the field, particularly the UK. Laboratory exposures also involved constant temperatures, which do not reflect the natural thermal fluctuations that can induce chill-hardening or allow partial recovery between cold events.

The long-term maintenance of laboratory colonies may also contribute to underestimation of cold tolerance. The *C. nubeculosus* colony used in this study has been reared at approximately 27 °C for over 50 years, during which time relaxed selection pressures may have led to the loss of heritable cold-tolerance traits and the development of culture-specific adaptations. Consequently, laboratory populations are likely less tolerant of low temperatures than wild conspecifics, meaning that cold tolerance observed under laboratory conditions may underestimate the survival potential of natural populations. However, differences between colony and field-caught larvae have been reported in *C. sonorensis*, where colony larvae were more cold-tolerant than field-caught larvae (McDermott et al. 2017). Similarly, survival recorded under controlled conditions may not fully represent ecological viability, as adult *Culicoides* activity in the field declines sharply below about 4 °C (Tugwell et al., 2021).

The capacity of *C. nubeculosus* to tolerate low temperatures across developmental stages may influence the potential for arbovirus persistence during winter. In temperate regions, viruses such as BTV and SBV can persist through a combination of vector, host, and environmental mechanisms (Wilson et al. 2008, Brand and Keeling 2017, Mansfield et al. 2024). The ability of larvae, pupae, and adults to survive temperatures below those typically experienced in northern Europe suggests that several developmental stages may remain viable through cold periods, most notably adults, that could support low-level viral persistence when transmission is otherwise limited. Experimental evidence indicates that BTV replication is highly dependent on temperature, with viral replication ceasing below approximately 12 °C (Carpenter et al. 2011, Brand and Keeling 2017). Potential overwintering mechanisms for BTV include transplacental transmission (which has been observed in the field), persistent viraemia in ruminant hosts, vertical transmission of virus from adult *Culicoides* to their offspring and long-lived infected adult *Culicoides* surviving through the winter. There has been no experimental evidence to support vertical transmission of BTV in *Culicoides,* but the cold tolerance observed in adult *Culicoides* in this study suggest that they may be able to survive substantial cold exposure, making this latter pathway a via option. This is particularly relevant when considering that adult *Culicoides* may move inside animal housing during the winter, where there is an abundance of hosts and warmer microclimates to aid survival. Previous studies have detected low activity levels of *Culicoides obsoletus* throughout the winter months indoors and outdoors in Germany (Groschupp et al. 2024). Furthermore, a study in Italy demonstrated that a higher proportion of *Culicoides* were found indoors when temperatures decreased outdoors (Magliano et al. 2018). Further research is required to determine the activity of adult *Culicoides* inside animal holdings within the UK.

Future studies should extend exposure durations to better reflect overwintering conditions, spanning weeks or months rather than hours to days. Incorporating fluctuating temperature regimes, rather than constant exposures, would more closely simulate natural winter conditions and reveal whether gradual cold cycles permit development at low temperatures. Experiments in natural substrates such as dung or saturated soil would add ecological realism by capturing the thermal buffering available in the field. Using field-caught midges across all life stages would provide essential context on how wild populations respond to cold stress. Findings from *C. sonorensis* indicate that cold tolerance can differ between populations, reinforcing the value of incorporating wild individuals into future life-stage comparisons (McDermott et al. 2017). Finally, following cold-exposed individuals through emergence, reproduction, and then through to the next generation would reveal whether sublethal or transgenerational effects of cold stress carry forward to the performance of the next generation; while pairing survival assays with physiological measurements could help identify the mechanisms underpinning cold tolerance.

## Conclusion

This study demonstrates that *C. nubeculosus* exhibits considerable cold tolerance across its life cycle, potentially enabling overwintering through multiple developmental stages rather than reliance on a single form. The ability of eggs and pupae to endure subzero temperatures, together with the short-term resilience of larvae and adults, suggests that populations can persist through transient cold events and recover when conditions improve. This multi-stage resilience may maintain vector populations between seasons and sustain arbovirus transmission potential through winter. Understanding how these overwintering mechanisms interact with environmental variability will provide valuable insight into the seasonal ecology, persistence, and transmission potential of *Culicoides* populations.

## Acknowledgments

The authors acknowledge Simon King for designing and constructing the specialised cage apparatus used to fully submerge experimental pots. We also recognise the Environmental Change Biology Laboratory.

## References

Alto BW, Reiskind MH, Lounibos LP. 2008. Size alters susceptibility of vectors to dengue virus infection and dissemination. Am J Trop Med Hyg 79(5):688–695. 10.4269/ajtmh.2008.79.688.

Banerjee P, Sarkar A, Harsha R et al. 2024. Influence of Rearing Temperatures on Oviposition and Survivability of Developmental Stages of *Culicoides peregrinus* Vector of Bluetongue Virus with a Note on Egg Viability. Proceedings of the Zoological Society 77(2):232–239. 10.1007/s12595-024-00527-3.

Bellekom B, Bailey A, England M et al. 2023. Effects of storage conditions and digestion time on DNA amplification of biting midge (*Culicoides*) blood meals. Parasites & Vectors 16(1). 10.1186/s13071-022-05607-x.

Bishop AL, McKenzie HJ. 1994. Overwintering of *Culicoides* spp. (Diptera: Ceratopogonidae) in the Hunter Valley, New South Wales. Australian Journal of Entomology 33(2):159163. 10.1111/j.1440-6055.1994.tb00943.x.

Blackwell A, King FC. 1997. The vertical distribution of *Culicoides impunctatus* larvae. Medical and Veterinary Entomology 11(1):45–48. 10.1111/j.1365-2915.1997.tb00288.x.

Boorman J. 1974. The maintenance of laboratory colonies of *Culicoides variipennis* (Coq.), *C. nubeculosus* (Mg.) and *C. riethi* Kieff. (Diptera, Ceratopogonidae). Bulletin of Entomological Research 64(3):371–377. 10.1017/S0007485300031254.

Brand SP, Keeling MJ. 2017. The impact of temperature changes on vector-borne disease transmission: *Culicoides* midges and bluetongue virus. J R Soc Interface 14(128). 10.1098/rsif.2016.0481.

Byttebier B, De Majo MS, Fischer S. 2014. Hatching Response of *Aedes aegypti (*Diptera: Culicidae) Eggs at Low Temperatures: Effects of Hatching Media and Storage Conditions. Journal of Medical Entomology 51(1):97–103. 10.1603/me13066.

Caminade C, McIntyre KM, Jones AE. 2019. Impact of recent and future climate change on vector-borne diseases. Ann N Y Acad Sci 1436(1):157–173. 10.1111/nyas.13950.

Carpenter S, Groschup MH, Garros C et al. 2013. *Culicoides* biting midges, arboviruses and public health in Europe. Antiviral Res 100(1):102–113. 10.1016/j.antiviral.2013.07.020.

Carpenter S, Wilson A, Barber J et al. 2011. Temperature Dependence of the Extrinsic Incubation Period of Orbiviruses in *Culicoides* Biting Midges. PLoS ONE 6(11):e27987. 10.1371/journal.pone.0027987.

Chala B, Hamde F. 2021. Emerging and Re-emerging Vector-Borne Infectious Diseases and the Challenges for Control: A Review. Frontiers in Public Health 9. 10.3389/fpubh.2021.715759.

Chapman S, Murphy E, Stainforth D et al. 2020. Trends in Winter Warm Spells in the Central England Temperature Record. Journal of Applied Meteorology and Climatology 59. 10.1175/JAMC-D-19-0267.1.

Chen X, Yan H, Ma L et al. 2024. Comprehensive experimental study of microbial respiration during self-heating in biomass storage piles. Fuel 362:130746. 10.1016/j.fuel.2023.130746.

Collins ÁB, Mee JF, Doherty ML et al. 2018. *Culicoides* species composition and abundance on Irish cattle farms: implications for arboviral disease transmission. Parasites & Vectors 11(1). 10.1186/s13071-018-3010-6.

Cribb BW, Chitra E. 1998. Ultrastructure of the eggs of *Culicoides molestus* (Diptera: Ceratopogonidae). J Am Mosq Control Assoc 14(4):363–368.

Devlin JJ, Unfried L, Lecheta MC et al. 2022. Simulated winter warming negatively impacts survival of Antarctica’s only endemic insect. Functional Ecology 36(8):1949–1960. 10.1111/1365-2435.14089.

Dijkstra E, Santman-Berends IMGA, Stegeman A et al. 2025. Evaluating the effect of vaccination on the impact of BTV-3 in Dutch sheep flocks in 2024. Veterinary Microbiology 309:110652. 10.1016/j.vetmic.2025.110652.

Ebrahimi B, Shakibi S, Foster WA. 2014. Delayed Egg Hatching of *Anopheles gambiae* (Diptera: Culicidae) Pending Water Agitation. Journal of Medical Entomology 51(3):580–590. 10.1603/me13100.

England ME, Pearce-Kelly P, Brugman VA et al. 2020. *Culicoides* species composition and molecular identification of host blood meals at two zoos in the UK. Parasites & Vectors 13(1). 10.1186/s13071-020-04018-0.

Erram D, Blosser EM, Burkett-Cadena N. 2019. Habitat associations of *Culicoides* species (Diptera: Ceratopogonidae) abundant on a commercial cervid farm in Florida, USA. Parasites & Vectors 12(1). 10.1186/s13071-019-3626-1.

Gołębiowski M, Boguś MI, Paszkiewicz M et al. 2011. Cuticular lipids of insects as potential biofungicides: methods of lipid composition analysis. Anal Bioanal Chem 399(9):3177–3191. 10.1007/s00216-010-4439-4.

Groschupp S, Kampen H, Werner D. 2024. Winter activity of *Culicoides* (Diptera: Ceratopogonidae) inside and outside stables in Germany. Medical and Veterinary Entomology 38(4):552–565. 10.1111/mve.12756.

Gubler DJ. 2010. The Global Threat of Emergent/Re-emergent Vector-Borne Diseases’. Springer Netherlands. 39-62.

Hadj-Henni L, Millot C, Lehrter V et al. 2023a. Wing morphometrics of biting midges (Diptera: *Culicoides*) of veterinary importance in Madagascar. Infection, Genetics and Evolution 114:105494. 10.1016/j.meegid.2023.105494.

Hadj-Henni L, Djerada Z, Millot C et al. 2023b. Wing morphology variations in *Culicoides circumscriptus* from France. Frontiers in Veterinary Science 10. 10.3389/fvets.2023.1089772.

Hochstrasser AL, Mathis A, Verhulst NO. 2024. Thermal preference of *Culicoides* biting midges in laboratory and semi-field settings. Journal of Thermal Biology 119:103783. 10.1016/j.jtherbio.2024.103783.

Juliano SA, Ribeiro GS, Maciel-De-Freitas R et al. 2014. She’s a femme fatale: low-density larval development produces good disease vectors. Memórias do Instituto Oswaldo Cruz 109(8):1070–1077. 10.1590/0074-02760140455.

Kameke D, Kampen H, Wacker A et al. 2021. Field studies on breeding sites of *Culicoides* Latreille (Diptera: Ceratopogonidae) in agriculturally used and natural habitats. Scientific Reports 11(1). 10.1038/s41598-021-86163-9.

King S, Nicholls M, Scales J et al. 2025. The efficacy of vector-proof accommodation for the protection of livestock against *Culicoides* biting midges. Parasites & Vectors 18(1). 10.1186/s13071-025-06736-9.

Koštál V, Grgac R, Korbelová J. 2019. Delayed mortality and sublethal effects of cold stress in *Drosophila melanogaster*. J Insect Physiol 113:24–32. 10.1016/j.jinsphys.2019.01.003.

Liao J, Liu J, Guan Z et al. 2021. Duration of Low Temperature Exposure Affects Egg Hatching of the Colorado Potato Beetle and Emergence of Overwintering Adults. Insects 12(7):609. 10.3390/insects12070609.

Lü J, Bai C, Guo Y et al. 2025. Influence of Cold Exposure for Different Durations on Laboratory-Reared *Habrobracon hebetor* (Say) (Hymenoptera: Braconidae). Diversity 17(4):253. 10.3390/d17040253.

Lühken R, Kiel E, Steinke S. 2014. *Culicoides* biting midge density in relation to the position and substrate temperature in a cattle dung heap. Parasitology Research 113(12):4659–4662. 10.1007/s00436-014-4182-4.

Lysyk TJ, Danyk T. 2007. Effect of Temperature on Life History Parameters of Adult *Culicoides sonorensis* (Diptera: Ceratopogonidae) in Relation to Geographic Origin and Vectorial Capacity for Bluetongue Virus. Journal of Medical Entomology 44(5):741–751. 10.1093/jmedent/44.5.741.

Machida WS, Tidon R, Klaczko J. 2022. Wing plastic response to temperature variation in two distantly related Neotropical *Drosophila* species (Diptera, Drosophilidae). Canadian Journal of Zoology 100(2):82–89. 10.1139/cjz-2021-0099.

MacMillan H, Sinclair B. 2010. Mechanisms underlying insect chill-coma. Journal of insect physiology 57:12–20. 10.1016/j.jinsphys.2010.10.004.

Macmillan HA, Findsen A, Pedersen TH et al. 2014. Cold-induced depolarization of insect muscle: Differing roles of extracellular K+ during acute and chronic chilling. Journal of Experimental Biology 217(16):2930–2938. 10.1242/jeb.107516.

Magliano A, Scaramozzino P, Ravagnan S et al. 2018. Indoor and outdoor winter activity of *Culicoides* biting midges, vectors of bluetongue virus, in Italy. Medical and Veterinary Entomology 32(1):70–77. 10.1111/mve.12260.

Mansfield KL, Schilling M, Sanders C et al. 2024. Arthropod-Borne Viruses of Human and Animal Importance: Overwintering in Temperate Regions of Europe during an Era of Climate Change. Microorganisms 12(7):1307. 10.3390/microorganisms12071307.

McDermott EG, Mullens BA. 2014. Desiccation Tolerance in the Eggs of the Primary North American Bluetongue Virus Vector, *Culicoides sonorensis* (Diptera: Ceratopogonidae), and Implications for Vector Persistence. J Med Entomol 51(6):1151–1158. 10.1603/me14049.

McDermott EG, Mayo CE, Mullens BA. 2017. Low Temperature Tolerance of *Culicoides sonorensis* (Diptera: Ceratopogonidae) Eggs, Larvae, and Pupae From Temperate and Subtropical Climates. J Med Entomol 54(2):264–274. 10.1093/jme/tjw190.

Modrý D, Hainisch E, Fuehrer H-P et al. 2025. Emergence of Autochthonous *Leishmania* (Mundinia) *martiniquensis* Infections in Horses, Czech Republic and Austria, 2019– 2023. Emerging Infectious Diseases 31. 10.3201/eid3109.250254.

Montini P, De Majo MS, Fischer S. 2021. Delayed mortality effects of cold fronts during the winter season on *Aedes aegypti* in a temperate region. J Therm Biol 95:102808. 10.1016/j.jtherbio.2020.102808.

Mullens BA, Rodriguez JL. 1992. Survival and Vertical Distribution of Larvae of *Culicoides variipennis* (Diptera: Ceratopogonidae) in Drying Mud Habitats. Journal of Medical Entomology 29(5):745–749. 10.1093/jmedent/29.5.745.

Neupane S, Davis T, Olds C et al. 2025. Unraveling the relationships between midge abundance and incidence, microbial communities, and soil and water properties in a protected natural tallgrass prairie. Parasites & Vectors 18(1). 10.1186/s13071-025-06780-5.

Nunamaker RA. 1993. Rapid Cold-Hardening in *Culicoides variipennis sonorensis* (Diptera: Ceratopogonidae). Journal of Medical Entomology 30(5):913–917. 10.1093/jmedent/30.5.913.

Ottih EC, Tripet F. 2024. Natural variation in timing of egg hatching, response to water agitation, and bidirectional selection of early and late hatching strains of the malaria mosquito *Anopheles gambiae sensu lato*. Parasites & Vectors 17(1). 10.1186/s13071-024-06533-w.

Overgaard J, Macmillan HA. 2017. The Integrative Physiology of Insect Chill Tolerance. Annual Review of Physiology 79(1):187–208. 10.1146/annurev-physiol022516-034142.

Purse BV, Carpenter S, Venter GJ et al. 2015. Bionomics of Temperate and Tropical *Culicoides* Midges: Knowledge Gaps and Consequences for Transmission of *Culicoides*-Borne Viruses. Annual Review of Entomology 60(1):373–392. 10.1146/annurev-ento-010814-020614.

Rozo-Lopez P, Park Y, Drolet BS. 2022. Effect of Constant Temperatures on *Culicoides sonorensis* Midge Physiology and Vesicular Stomatitis Virus Infection. Insects 13(4):372.

Ryan SJ, Carlson CJ, Mordecai EA et al. 2019. Global expansion and redistribution of *Aedes*-borne virus transmission risk with climate change. PLOS Neglected Tropical Diseases 13(3):e0007213. 10.1371/journal.pntd.0007213.

Sanders CJ, Shortall CR, England M et al. 2019. Long-term shifts in the seasonal abundance of adult *Culicoides* biting midges and their impact on potential arbovirus outbreaks. Journal of Applied Ecology 56(7):1649–1660. 10.1111/13652664.13415.

Schneider CA, Rasband WS, Eliceiri KW. 2012. NIH Image to ImageJ: 25 years of image analysis. Nature Methods 9(7):671–675. 10.1038/nmeth.2089.

Sherpa S, Tutagata J, Gaude T et al. 2022. Genomic Shifts, Phenotypic Clines, and Fitness Costs Associated With Cold Tolerance in the Asian Tiger Mosquito. Molecular Biology and Evolution 39(5). 10.1093/molbev/msac104.

Sick F, Beer M, Kampen H et al. 2019. *Culicoides* Biting Midges—Underestimated Vectors for Arboviruses of Public Health and Veterinary Importance. Viruses 11(4):376. 10.3390/v11040376.

Sinclair BJ, Ferguson LV, Salehipour-Shirazi G et al. 2013. Cross-tolerance and Cross-talk in the Cold: Relating Low Temperatures to Desiccation and Immune Stress in Insects. Integrative and Comparative Biology 53(4):545–556. 10.1093/icb/ict004.

Slama D, Baraket R, Remadi L et al. 2021. Morphological and molecular differentiation between *Culicoides oxystoma* and *Culicoides kingi* (Diptera: Ceratopogonidae) in Tunisia. Parasites & Vectors 14(1):607. 10.1186/s13071-021-05084-8.

Socha W, Kwasnik M, Larska M et al. 2022. Vector-Borne Viral Diseases as a Current Threat for Human and Animal Health—One Health Perspective. Journal of Clinical Medicine 11(11):3026. 10.3390/jcm11113026.

Steinke S, Lühken R, Kiel E. 2014. Assessment of the abundance of *Culicoides chiopterus* and *Culicoides dewulfi* in bovine dung: A comparison of larvae extraction techniques and emergence traps. Veterinary Parasitology 205(1):255–262. 10.1016/j.vetpar.2014.07.030.

Steyn J, Venter GJ, Labuschagne K et al. 2016. Possible over-wintering of bluetongue virus in *Culicoides* populations in the Onderstepoort area, Gauteng, South Africa. Journal of the South African Veterinary Association 87(1). 10.4102/jsava.v87i1.1371.

Stokes JE, Carpenter S, Sanders C et al. 2022. Emergence dynamics of adult *Culicoides* biting midges at two farms in south-east England. Parasites & Vectors 15(1). 10.1186/s13071-022-05370-z.

Terhzaz S, Teets NM, Cabrero P et al. 2015. Insect capa neuropeptides impact desiccation and cold tolerance. Proc Natl Acad Sci U S A 112(9):2882–2887. 10.1073/pnas.1501518112.

Thompson GM, Jess S, Murchie AK. 2013. Differential emergence of *Culicoides* (Diptera: Ceratopogonidae) from on-farm breeding substrates in Northern Ireland. Parasitology 140(6):699–708. 10.1017/s0031182012002016.

Tugwell LA, England ME, Gubbins S et al. 2021. Thermal limits for flight activity of field-collected *Culicoides* in the United Kingdom defined under laboratory conditions. Parasites & Vectors 14(1). 10.1186/s13071-020-04552-x.

Uslu U, Dik B. 2006. Vertical distribution of *Culicoides* larvae and pupae. Medical and Veterinary Entomology 20(3):350–352. 10.1111/j.13652915.2006.00626.x.

Uslu U, Dik B. 2010. Chemical characteristics of breeding sites of *Culicoides* species (Diptera: Ceratopogonidae). Veterinary Parasitology 169(1):178–184. 10.1016/j.vetpar.2009.12.007.

Van Den Eynde C, Sohier C, Matthijs S et al. 2021. Temperature and food sources influence subadult development and blood-feeding response of *Culicoides obsoletus (sensu lato)* under laboratory conditions. Parasites & Vectors 14(1). 10.1186/s13071-021-04781-8.

Werner D, Groschupp S, Bauer C et al. 2020. Breeding Habitat Preferences of Major *Culicoides* Species (Diptera: Ceratopogonidae) in Germany. International Journal of Environmental Research and Public Health 17(14):5000. 10.3390/ijerph17145000.

White DM, Wilson WC, Blair CD et al. 2005. Studies on overwintering of bluetongue viruses in insects. J Gen Virol 86(Pt 2):453–462. 10.1099/vir.0.80290-0.

White SM, Sanders CJ, Shortall CR et al. 2017. Mechanistic model for predicting the seasonal abundance of *Culicoides* biting midges and the impacts of insecticide control. Parasites & Vectors 10(1). 10.1186/s13071-017-2097-5.

Wilson A, Darpel K, Mellor PS. 2008. Where Does Bluetongue Virus Sleep in the Winter? PLoS Biology 6(8):e210. 10.1371/journal.pbio.0060210.

Wu S, Luo M, Lau GN-C et al. 2025. Rapid flips between warm and cold extremes in a warming world. Nature Communications 16(1). 10.1038/s41467-02558544-5.

