## Supplementary Material for "Substantial cold tolerance in all life stages of the biting midge, *Culicoides nubeculosus* (Diptera: Ceratopogonidae)"

**Table S1.** Summary of the container setup and substrate composition used for each *Culicoides nubeculosus* life stage during temperature exposure experiments.

| **Life stage** | **Container setup** | **Substrate and medium** |
| --- | --- | --- |
| **Eggs** | 35mm petri dish inside 125ml pot | Polyester wadding with 3ml rearing fluid + filter paper |
| **L1** | 125ml pot | Polyester wadding with 15ml rearing fluid |
| **L4** | 125ml pot | Polyester wadding with 15ml rearing fluid |
| **Pupae** | 125ml pot | Polyester wadding with 12ml rearing fluid + filter paper |
| **Adults** | 125ml pot | Dry filter paper |

**Table S2.**  Summary of statistical models used to assess the effects of low-temperature exposure on survival, development, emergence, and wing length across life stages of *Culicoides nubeculosus*. Each row (Models A–R) represents a distinct analysis conducted for a specific life stage and response variable. The table lists the model type (GLM, GLMM, or LMM), fixed and random effects included, whether control treatments were incorporated, how temperature was treated (categorical or continuous), and the type of analysis of deviance (ANOVA) test used. Firth’s bias correction was applied in binomial models where data separation occurred. Type III ANOVA tests were used for all models to evaluate fixed effects. Gaussian LMMs were used for continuous responses (wing length), and binomial or quasibinomial models were applied for proportional outcomes (survival, emergence, development success).

| **Model** | **Life Stage** | **Response** | **Model Type** | **Fixed Effects** | **Random Effects** | **Control Included** | **Temperature Type** | **ANOVA Test** |
| --- | --- | --- | --- | --- | --- | --- | --- | --- |
| A | All stages | Survival (assessed at life-stage specific endpoints) | GLM (quasibinomial) | Temperature * Life stage | None | No | Continuous | Type III (F test) |
| B | Egg | Hatch success | GLM (binomial) | Temperature * Duration | None | Yes | Categorical | Type III |
| C | Egg | Adult emergence | GLM (binomial, Firth) | Temperature * Duration | None | Yes | Categorical | Type III |
| D | Egg | Hatch rate (1H exposure) | GLMM (binomial) | Temperature * Day | Replicate | Yes | Categorical | Type III |
| E | Egg | Hatch Rate (1H,6H and 24H exposure) | GLMM (binomial) | Temperature * Duration * Day | Replicate | Yes | Categorical | Type III |
| F | L1 | Survival (28D endpoint) | GLM (quasibinomial) | Temperature * Duration | None | Yes | Categorical | Type III (F test) |
| G | L1 | Adult emergence | GLM (binomial, Firth) | Temperature + Duration | None | Yes | Categorical | Type III |
| H | L4 | Survival (1H post exposure) | GLM (binomial, Firth) | Temperature * Duration | None | Yes | Categorical | Type III |
| I | L4 | Survival (10D endpoint) | GLM (binomial, Firth) | Temperature * Duration | None | Yes | Categorical | Type III |
| J | L4 | Adult emergence | GLM (binomial, Firth) | Temperature + Duration | None | Yes | Categorical | Type III |
| K | Pupae | Emergence | GLM (binomial,  Firth) | Temperature * Duration | None | Yes | Categorical | Type III |
| L | Adult | Survival (1H post exposure) | GLM  (binomial) | Temperature * Duration | None | Yes | Categorical | Type III |
| M | Adult | Survival (3D endpoint) | GLM (binomial, Firth) | Temperature * Duration | None | Yes | Categorical | Type III |
| N | All stages | Wing size | LMM (Gaussian) | Temperature * Life stage + Sex | Replicate | No | Continuous | Type III |
| O | Eggs | Wing size | LMM (Gaussian) | Temperature + Duration + Sex | Replicate | Yes | Categorical | Type III |
| P | L1 | Wing size | LMM (Gaussian) | Temperature + Duration + Sex | Replicate | Yes | Categorical | Type III |
| Q | L4 | Wing size | LMM (Gaussian) | Temperature + Duration + Sex | Replicate | Yes | Categorical | Type III |
| R | Pupae | Wing size | LMM (Gaussian) | Temperature + Duration + Sex | Replicate | Yes | Categorical | Type III |

**Section S3. Developmental outcomes across life stages**

**Egg Developmental outcomes**

The proportion of individuals that reached adulthood after 28 days was significantly affected by the interaction between temperature and exposure duration (Table S2). At 1 h, adult emergence was higher in the control than at −11 °C (p < 0.001) and −14 °C (p = 0.004), with no other significant differences. At 6 h, emergence was higher at −1 °C than at −11 °C (p = 0.031) and −14 °C (p < 0.001), and higher at −11 °C than at −14 °C (p = 0.019). Emergence at −11 °C (p = 0.010) and −14 °C (p < 0.001) was also lower than the control. At 24 h, emergence was lower at −1 °C than at −11 °C (p = 0.021) and the control (p = 0.016), and emergence at −14 °C was lower than both −11 °C (p = 0.005) and the control (p < 0.001). When temperature groups were pooled, exposure duration had no significant effect on adult emergence (Figure S1).

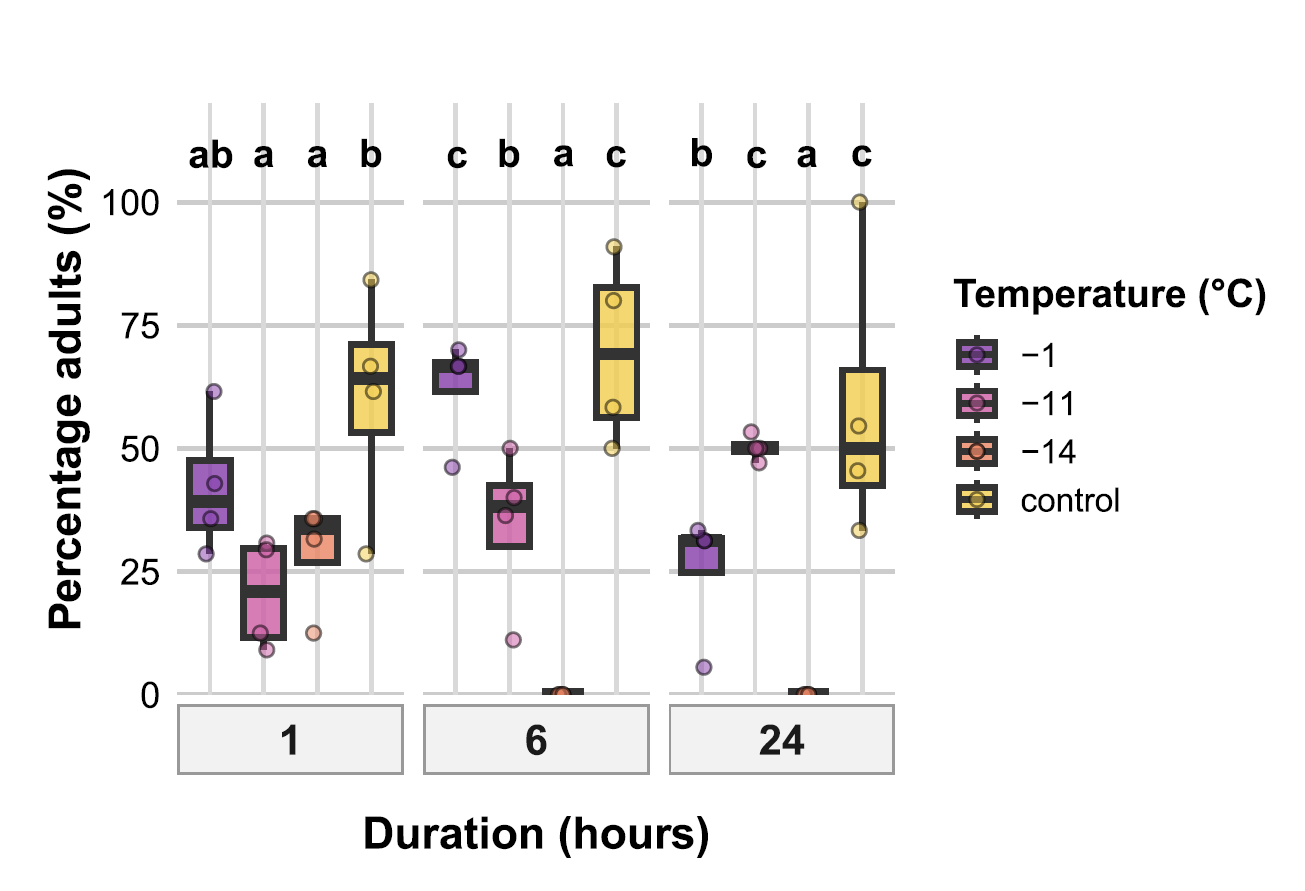

**Figure S1.** Percentage of *Culicoides nubeculosus* eggs that developed to adults after 28 days at different temperatures (−1, −11, −14 °C, and control) and exposure durations (1, 6, and 24 h). At 1 h, adult emergence was higher in the control than at −11 °C and −14 °C; at 6 h, emergence was higher at −1 °C than at −11 °C; and at 24 h, emergence was lower at −1 °C than at −11 °C and the control. Different letters indicate statistically significant differences among treatments (Tukey-adjusted *p* < 0.05).

**Alt text:** Graph showing the percentage of *Culicoides nubeculosus* eggs that developed into adults after 28 days at temperatures of −1, −11, −14 °C, and control and exposure durations of 1, 6 and 24 hours.

**Larval instar 1 (L1) developmental outcomes**

The proportion of L1 larvae that developed to adults was significantly affected by temperature but not by exposure duration (Table S2). Because several treatment combinations had low or no emergence, the interaction term was excluded. Adult emergence at −6 °C was significantly lower than at −1 °C (*p* < 0.001) and the control (*p* < 0.001). Emergence at −10 °C did not differ significantly from the other treatments. Duration effects were not statistically significant (Figure S1).

**L4 developmental outcomes**

The proportion of adults among surviving L4 larvae was significantly affected by both temperature and exposure duration (Table S2). Because some treatment combinations (−7 °C at 6 h and 24 h) had no survivors, the interaction term was excluded, and an additive bias-reduced binomial model was used. Adult emergence at −6 °C was significantly lower than at −1 °C and the control (both *p* < 0.001), and emergence at −7 °C was also significantly lower than the control (*p* = 0.019). Differences between −1 °C and the control, and among exposure durations, were not statistically significant. (Figure S1).

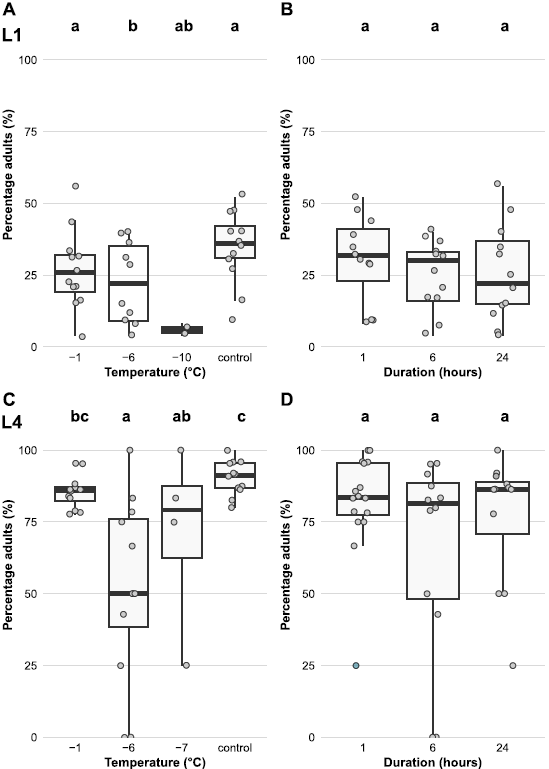

**Figure S2.** Percentage of *Culicoides nubeculosus* larvae that developed to adults for L1 (A–B) and L4 (C–D) following cold exposure. Panels show effects of temperature (A, C) and exposure duration (B, D). Only main effects were tested because some treatment combinations had zero or near-zero emergence, making the interaction term non-estimable. Different letters indicate statistically significant differences among treatments (Tukey-adjusted *p* < 0.05).

**Alt text:** Four graphs, the first showing the percentage of L1 larval *Culicoides nubeculosus* that successfully developed into adults at temperatures of −1, −6, −10 °C, and control; the second showing the percentage of L1 larvae that developed into adults after cold exposure durations of 1, 6 and 24 hours; the third showing the percentage of L4 larval *Culicoides nubeculosus* that successfully developed into adults at temperatures of −1, −6, −7 °C, and control; and the fourth showing the percentage of L4 larvae that developed into adults after cold exposure durations of 1, 6 and 24 hours.

**Table S3.** Analysis of deviance from binomial generalised linear models (GLMs) assessing the effects of temperature and exposure duration on developmental outcomes across egg, L1 larval, and L4 larval stages of *Culicoides nubeculosus*. For eggs, a full model including the temperature × duration interaction was fitted. For L1 and L4 larvae, additive bias-reduced binomial models were used due to low or zero survival in some treatment combinations. Reported values are likelihood ratio chi-squared (χ²) statistics with associated degrees of freedom (df) and p-values from Type III analyses of deviance.

| **Developmental Stage** | **Factor** | **Chi-squared** | **df** | **P value** |
| --- | --- | --- | --- | --- |
| **Eggs** | Temperature | 138.282 | 3 | **< 0.001** |
|  | Duration (hours) | 11.442 | 2 | **0.003** |
|  | Temperature × Duration | 68.73 | 6 | **< 0.001** |
| **L1 Larvae** | Temperature | 31.320 | 3 | **< 0.001** |
|  | Duration (hours) | 2.979 | 2 | 0.226 |
| **L4 Larvae** | Temperature | 40.721 | 3 | **< 0.001** |
|  | Duration (hours) | 6.731 | 2 | **0.035** |
